# Testing the adaptive value of sporulation in budding yeast using experimental evolution

**DOI:** 10.1101/2020.02.23.959684

**Authors:** Kelly M. Thomasson, Alexander Franks, Henrique Teotónio, Stephen R. Proulx

## Abstract

*Saccharomyces* yeast can grow through mitotic vegetative cell division while they convert resources in their environment into biomass. When cells encounter specific low nutrient environments, sporulation may be initiated and meiotic division produces 4 haploid cells contained inside a protective ascus. The protected spore state does not acquire new resources but is partially protected from desiccation, heat, and caustic chemicals. Because cells cannot both be protected and acquire resources simultaneously, committing to sporulation represents a trade-off between current and future reproduction. Recent work has suggested that one of the major environmental factors that select for the formation of spores is passaging through insect guts, as this also represents a major way that yeasts are vectored to new food sources. We subjected replicate populations of a panel of 5 yeast strains to repeated, predictable passaging through insects by feeding them to fruit flies (*Drosopila melanogaster*) and then allowing surviving yeast cell growth in defined media for a fixed period of time. We also evolved control populations using the same predictable growth environments but without being exposed to flies. We assayed populations for their sporulation rate, as measured by the percentage of cells that had sporulated after resource depletion. We found that the strains varied in their ancestral sporulation rate such that domesticated strains had lower sporulation. During evolution, all strains evolved increased sporulation in response to passaging through flies, but domesticated strains evolved to lower final levels of sporulation. We also found that exposure to flies led to an evolved change in the timing of the sporulation response relative to controls, with a more rapid shift to sporulation, and that wild-derived strains showed a more extreme response. We conclude that strains that have lost the ability to access genetic variation for total sporulation rate and the ability to respond to cues in the environment that favor sporulation due to genetic canalization during domestication.

## Introduction

Most organisms have specialized life-history stages for growth when environmental conditions are favorable or life-history stages for dispersal to novel habitats when environmental conditions become challenging. Evolutionary theory predicts that natural selection will favor genotypes that maximize the relative fitness of expressing these life-history transitions as a function of the predictability of environmental change and the spatial structure of the populations in question (Olivieri *et al*., 1995; Tufto, 2000). Alternatively, fitness trade-offs between life-history stages can be due to genetic constraints between the relevant growth and dispersal traits, either because of negative genetic correlations if evolution occurs from standing genetic variation or lack of *de novo* mutational options (Lande, 1980).

In nature, populations of the budding yeast *Saccharomyces cerevisiae* are thought to primarily grow as a vegetative mitotic diploid cells and to disperse to novel habitats through the guts of insect vectors as meiotic haploid quiescent spores encapsulated within a protective structure called the ascus (Stefanini *et al*., 2012; Gibbs and Stanton, 2001; Coluccio *et al*., 2008). Sporulation in turn is initiated when diploid cells encounter adverse environments (Neiman, 2005). Several strains of *S. cerevisiae* have been domesticated and in these sporulation can be induced by changes in nutrient availability and pH, thought to be correlated with changes in resource availability that indicate starvation (Neiman, 2005). Domesticated strains appear however to have lost much of their ancestral sporulation efficiency (as measured by sporulation rates), when compared to wild strains (Gerke *et al*., 2006; De Chiara *et al*., 2020), presumably because they have been cultured as growing vegetative cells for many generations and maintaining the potential to sporulate is developmentally and physiologically costly when dispersal is no longer assured by insects (Ratcliff *et al*., 2013). Spores and vegetative cells can both survive insect guts, although with different success rates (Coluccio *et al*., 2008), and only spores are be able to cross-fertilize spores from other asci which can increase genetic variation available for future adaptation to local environmental conditions (Reuter *et al*., 2007).

Explanations about the evolution of sporulation have focused on a scenario where populations periodically experience ingestion by insects that cause high mortality in vegetative cells while allowing survival of spores (Ratcliff *et al*., 2013; Coluccio *et al*., 2008). In this scenario, the fitness of a genotype depends on how many progeny cells survive ingestion and establish new colonies in the next habitat when the insect defecates. If ingestion by insects happens only rarely, or only after the local resources are depleted, then we expect genotypes that maximally convert resources into spores will evolve. In these circumstances, selection should favor genotypes that sporulate only after most nutrients are depleted and vegetative diploids experience starvation conditions. Ignoring group-level traits (but see Discussion), individual *S. cerevisiae* growth and fitness depends both on traits that confer local adaptation and traits that allow cells to accurately sense unfavorable environmental conditions and only then to initiate and complete sporulation. On the other hand, *S. cerevisiae* dispersal ability depends on the ability to survive insect guts (as spores or vegetative cells) as well as mate recognition and germination once a new suitable growth habitat is reached (Murphy and Zeyl, 2012).

We set out to test if fitness-trade offs between growth and dispersal traits in *S. cerevisiae* are genetically constrained. For this we performed a replicated selection experiment in five *S. cerevisiae* strains that varied in their level of domestication, and where cultures were given time to exhaust their nutrient and were then propagated either through the guts of *Drosophila melanogaster* fruit-flies or directly transferred. All strains were initially isogenic and homothalic, such that evolution during the experiment could only occur through selection on *de novo* mutations and with little opportunity for recombination during the experiment. If fitness trade-offs result from a genetic constraint, then domesticated strains may have lost most of their ability to respond to selection for increased or faster sporulation, when compared with the wild strains. An alternative mode of adaptation to insect passaging would be to evolve increased vegetative cell survival through the *D. melanogaster* gut. The selection response of populations that were directly transferred allow us to disentangle genetic constraints between sensing starvation and initiation of sporulation independently of *D. melanogaster* gut vectoring.

## Methods

### Yeast and fruit-fly strains

We used a set of five genetically distinct strains of *S. cerevisiae* that were provided as homozygous diploids with resistance to Geneticin (G418), an orthologue to kanamycin, (Louvel *et al*., 2014) obtained from the National Collection of Yeast Cultures, UK. These five strains represent a wide range of ecological backgrounds including an Oak woodland in the Northeast united states (North American strain, AM), a palm flower in a Malaysian forest (Malaysian, MY), a West African strain from a semi-natural beer fermentation (West African, WA), a Sake brewery in Japan (Japanese Sake, JS), and a winery in Western Europe (Wine European, WE). These strains cover a range of backgrounds from wild, to partially human associated, to fully domesticated. Each strain was genetially modified for use in a lab setting by knocking out the mating type switching locus and adding DNA barcodes with stable diploid strains produced by complementary mating.

Each Ancestor was split into 4 replicate populations, which were then split into control and treatment populations. These 40 experimental populations were evolved for 30 full cycles of population growth, starvation, and passaging. Passaging was through Drosophila feeding in the treatment populations and by pipette for the control populations.

The *D. melanogaster* stocks used in one of the selection treatments (see next section) were created by outcrossing strains from isogenic Al-Ral, Taiwanese, Santa Barbarian and Malaysian lines. Flies to be used as vectors during the experiments were allowed to lay eggs on YPD agar plates. Adult flies were then removed and the eggs were bleached using a 10% bleach solution for 40 minutes at 22 °C. Fly eggs were collected by sterile pipette, washed with sterile water, and transferred using sterile technique to clean media. Clean flies were reared and propagated on this media so that other yeasts and fungi were minimized, but also so that the ingestion of antifungal elements did not reduce the viability of living yeasts traveling through the gut.

### Selection protocol

We grew each of our five diploid yeast strains in 2 mL of YPD (Yeast-Peptone-Dextrose) liquid culture over a 5 day period. Samples of these initial strains were then frozen in 15% glycerol solution at −80 °C and labelled as the “Ancestral” treatment. The five strains were split into 4 replicate each, and each was then split again into a paired control and treatment population (Figure 1).

**Figure 1:**
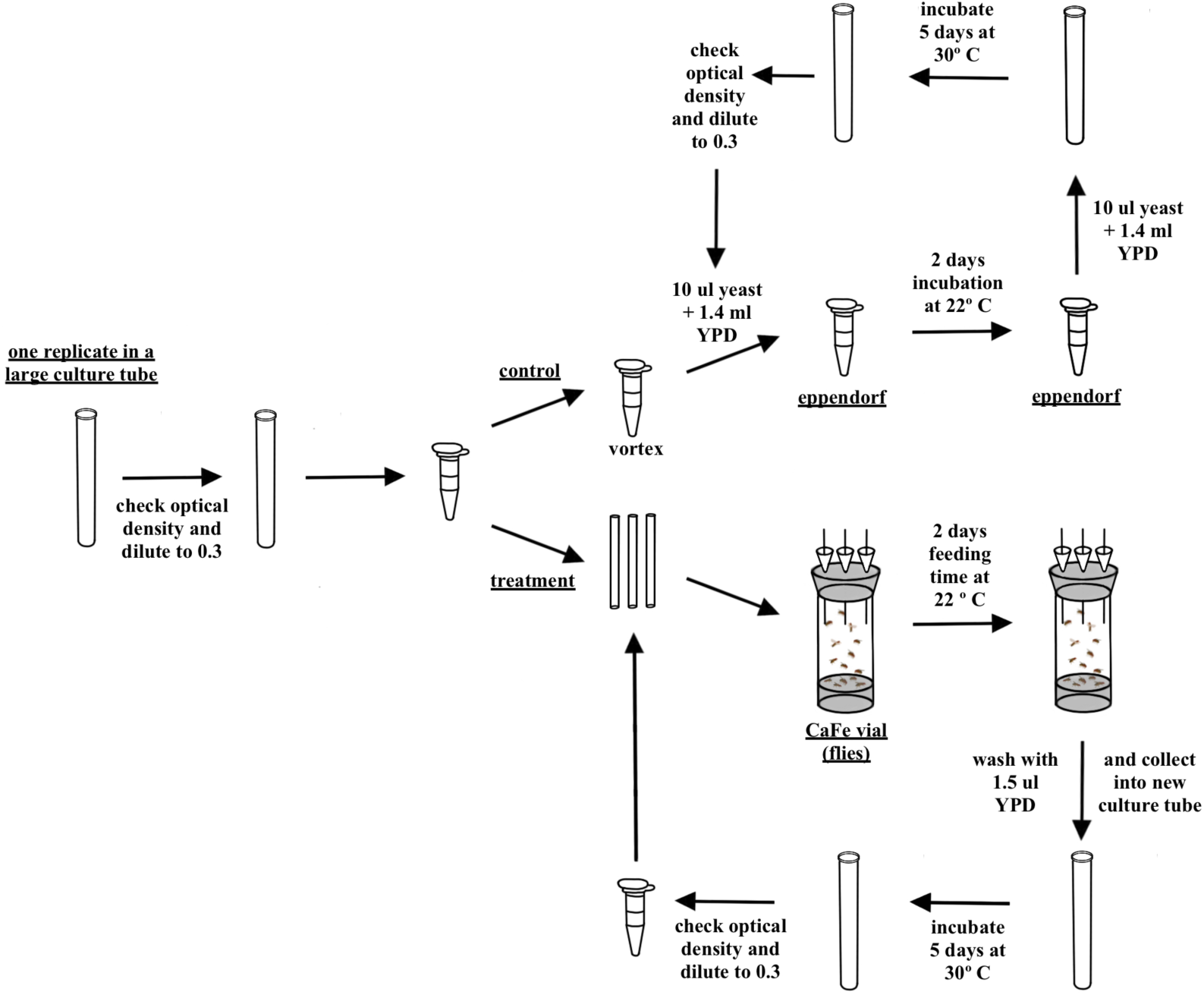
Each lineage was grown in liquid media and the sample was adjusted to an optical density of 0.3 and split into control and treatment tubes. From that point, the control and treatment went through parallel procedures lasting 7 days: 2 days of exposure to treatment (control: 22 °C bench-top incubation; treatment: 22 °C exposure to flies) and 5 days of growth or sporulation in liquid media at 30 °C. This was repeated for 30 cycles.

At the start of each selection cycle, each population was incubated in YPD at 30°C for 5 days without shaking. In order to reduce chances of contamination from other yeasts or bacteria, cultures were always grown in YPD media with G418, tetracycline, and ampicillin added. We performed experiments without shaking to increase the opportunities for haploid cells developing from spores to mate and form diploids. Under these growth conditions, yeast populations usually consume most of the sugar resources within 24 hours, and the 5 day period allowed ample time for spore formation to be initiated and completed.

After the 5 day period, each population was adjusted to an optical density of 0.3 in an effort to both ensure adequate population size and to prevent blockage in the capillary tube in the CaFe vial caused by high cell density (see below). The dilution process was performed using spent YPD, YPD that had been depleted by culturing yeast for 2 weeks and then filter sterilized. Spent YPD (SYPD) was used to mimic carbon and nitrogen sources in a late growth stage yeast culture population and to minimize population growth or germination of sporulated yeast cells (Madhani, 2007; Neiman, 2005).

For the fruit-fly experimental treatment we employed the Fly Capillary Feeder (CaFe) apparatus, adapted from Ja *et al.* (2007). CaFe tops used four 200*µ*L pipette tips which were cut to increase opening size, rubber stoppers and standard rubber bands. These CaFe tops were then affixed to narrow fly vials each containing 2ml of 3% solidified agarose solution to maintain humidity within the vial. The pipette tips within the CaFe top were fitted to one 5*µ*L capillary tube each. 18 hours prior to treatment, four clean, sexed flies were added to each of four vials per replicate. This starvation period was added to ensure sufficient consumption of yeast by the flies during treatment (Reuter *et al*., 2007).

Yeast cultures were distributed into CaFe feeding apparatus and offered to 3-4 clean flies (see above). Flies were allowed to feed for 48 hours and then removed from the vials. Measurements of total fly food consumption were taken by recording the change in meniscus of the two capillary tubes. Flies were removed from the vials which were then rinsed with 1.6 mL YPDA media and the supernatant (1.5 mL total volume because some volume is reabsorbed into the agar in the vial) was collected and used as the inoculate for the next round of yeast population growth.

For the control selection treatment yeast cultures were not distributed into the CaFe feeding apparatus, being instead placed on the bench-top nearby. After 48h, each control population was vortexed and 10*µ*L of each culture was moved to 1.49 mL of YPDA in a new culture tube (1.5 mL total volume).

These selection protocols were repeated for 30 growth cycles. Samples of each population were frozen every other cycle as a backup in case of contamination. After the last cycle all population were frozen and stored at −80 °C.

### Sporulation assay

The sporulation assay was performed in two blocks defined by the date of assay experiment, with each block containing samples of all 40 control and treatment replicate populations as well as the ancestral populations. Each ancestral population was assayed 6 times. In each block, two samples were taken from each population and then further divided into two technical replicates each for measurement. For the ancestral populations, three samples were taken in each assay block, also divided into two technical replicates.

We first grew thawed population samples on YPD agar plates and then inoculated liquid YPD cultures which grew for 6 hours without shaking at 30°C. Each sample was diluted to an optical density of 1.5 and 2 mL was centrifuged and washed to remove any traces of growth media. Cells were then resuspended in in 2 mL of Potassium Acetate (2% KAc at pH *≈* 6.7), and incubated at 30°C with shaking (230 rpm). Measurements were taken at 2.5 days and at 5 days. Samples were diluted by taking 5*µ*L of sample and 95*µ* L water. A field of cells was then photographed at 400*X* magnification. Images were processed by adding a grid and counting the number of spores and vegetative cells.

### Statistical analysis

We developed hierarchical models of sporulated cell counts where the observed counts were taken to be binomially distributed. We modeled the logit of the binomial parameter as a linear function of the interaction between selection treatment, and assay time along with a random effect of the replicate population, where we pooled variation among sampling and technical replicates. We performed these analyses separately on the Ancestral populations because they necessarily do not have the same structure of replicate experimental populations. For the experimentally evolved populations, we modelled each strain separately and then performed comparative analyses on the inferred posterior distributions.

The main model considers the sporulation probability to be a function of the treatment type (fly treatment or control) and the assay time (2.5 days or 5 days) giving

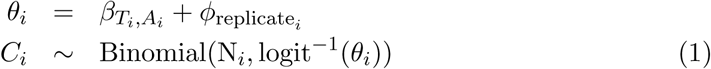

where *C*_*i*_ is the count of sporulated cells in sample *i*, N_*i*_ is the total number of cells counted in sample *i*. The predictor *β* depends on the *T* treatment type and the *A* assay time of observation *i*.

We additionally modeled the variation that comes from replicate populations sharing the same ancestry as

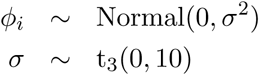

meaning that each replicate population is considered to be sampled from a normal distribution and the prior on the standard deviation is a Student’s t distribution with 3 degrees of freedom, a location of 0 and scale of 10.

Here, because we assume that the parameter for each combination of strain, treatment, and assay time is independent, this is similar to a generalized linear model with main effects and interaction terms for each of the three factors. The additional term *φ* represents the random effect of the four replicate populations and effectively allows for overdispersion of the count data relative to a binomial model. Such overdispersion in the count data could arise due to idiosyncratic differences during the creation of the replicate experimental populations from the ancestral strain.

We took a Bayesian approach for parameter inference and compared posterior distributions of parameter estimates as a way to evaluate hypotheses regarding the causes of selection responses. We specified the models using `BRMS` (Bürkner, 2017, 2018) and `STAN` (Stan Development Team, 2018) using `RStan` which performs Bayesian inference using a Hamiltonian Monte Carlo sampling to calculate the posterior probability of the model parameters given the observed data (`R` version 3.3.2, `RStan` version 2.15.1)(R Core Team, 2019). Convergence of the MCMC chains was checked by ensuring that 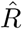 was less than 1.1, where 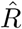 is a metric describing the variability between chains (Gelman *et al*., 2013).

### Data and code archiving

All data and code for analysis will be submitted to Data Dryad. Complete code to perform the analysis is available in a supplemental RMarkdown file.

## Results

### Ancestral sporulation rates

Sporulation rates at both assay times differ among the ancestral strains (Figure 2). We also compared the model with factorial effects of strain and assay time to a model with no effect of strain and found that the full model was highly supported (see supplementary rMarkdown file). As expected, this reflects their evolutionary history, with strains coming from natural environments showing higher sporulation rates while strains from industrial alcohol production showing lower sporulation rates (Gerke *et al*., 2006; De Chiara *et al*., 2020). The African strain WA, which is thought to be only partially-domesticated (see Methods) shows intermediate sporulation rates.

**Figure 2:**
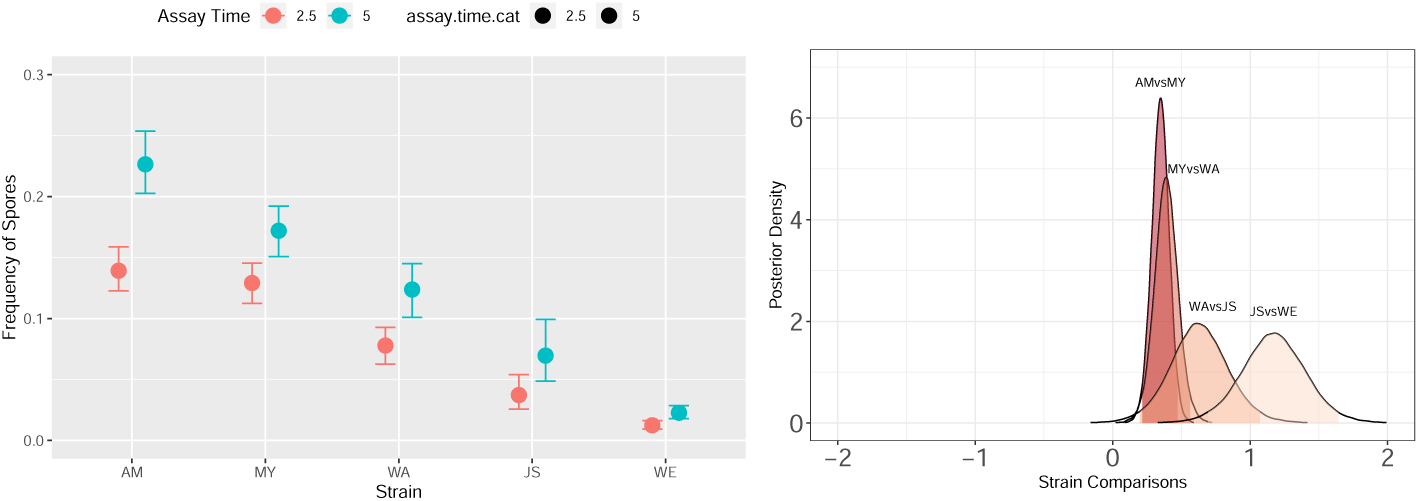
Sporulation rates in the ancestral strains. Panel A shows the fitted values for sporulation rate measured at 2.5 and 5 days (see equation 1), with the solid circles showing the median posterior value and the bars are the 95% posterior credible interval. The strains are arranged by decreasing ancestral sporulation rate, which also reflects the the degree of domestication (AM and MY, wild; WA, partially-domesticated; JS and WE, industrial). The difference between sporulation rate at 2.5 and 5 days can be interpreted as the speed at which vegetative cells enter and complete sporulation. Wild strains therefore show a faster transition to sporulation than domesticated strains. Panel B shows the pairwise comparisons between successive strains at the 5 day assay time point. The posterior probability of the difference being less than 0 is less than 0.001 for all comparisons

The sporulation rates of the strains are distinct as is the transition between days 2.5 and day 5, inferred by comparing models that do not include the strain or day of assay effect (see supplementary R markdown file). The strain effect can also be visualized by comparing the posterior distributions of the day five assay in a pairwise fashion between each successive strain to show that each strain has a distinct sporulation rate (figure 2)

The difference between sporulation rate at 2.5 and 5 days represents the speed at which vegetative cells enter and complete sporulation once they sense starvation conditions while in the culture tubes (Figure 1). For this trait, wild strains show a sharper change in completed sporulation between the time points as compared to domesticated strains (Figure 2). Note also that the strains are inferred to have distinct sporulation rates, as the posterior distributions between strains do not overlap at the 95% level.

### Selection responses in the fruit-fly treatment

After 30 growth cycles involving fruit-fly gut dispersal the evolved populations showed higher sporulation rates (Figure 3 and figure 4). For all strains, the posterior distribution is well away from 0 (the posterior probability of the difference from the ancestral being less than zero is less than 10^−3^), with sporulation rates of the evolved populations at the assay time of day 5 ranging from 30% to 95%. However, the magnitude of this effect is independent of the degree of domestication: for example, one of the wild strains (AM) shows the maximum response relative to its ancestral state, while another wild strain (MY) shows an intermediate response.

**Figure 3:**
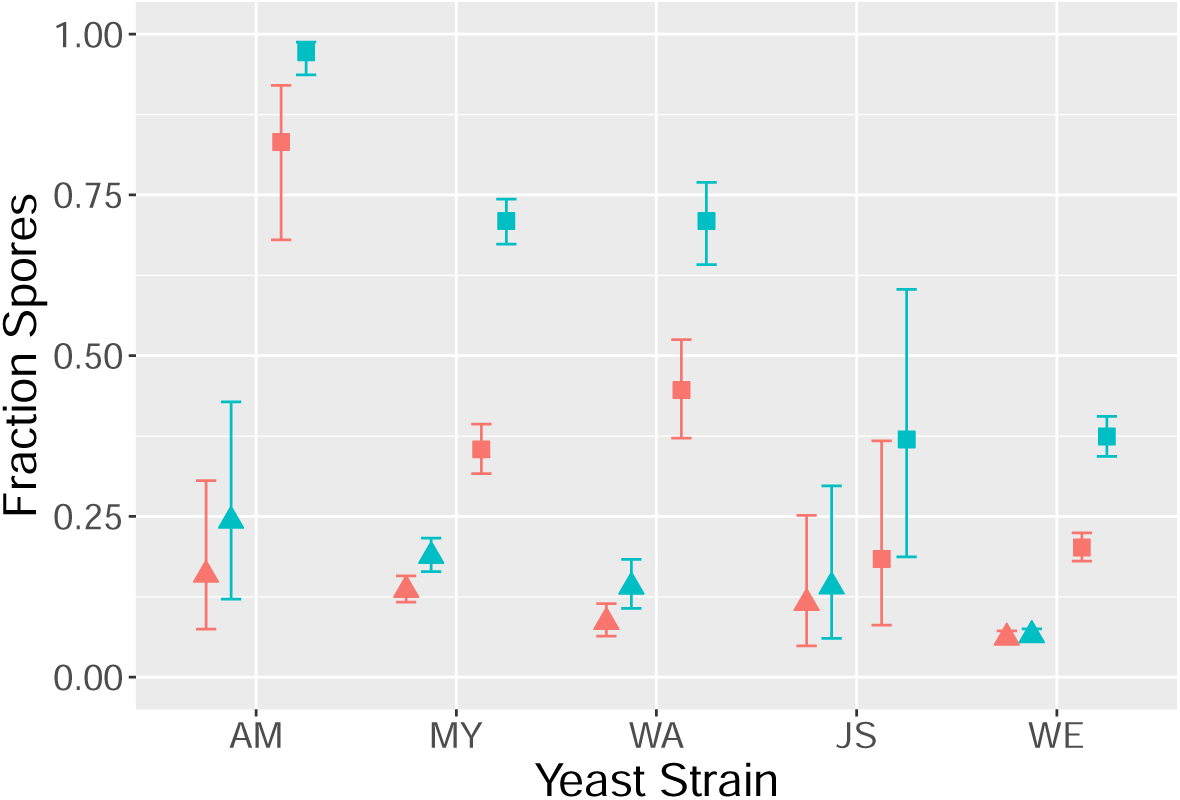
Inferred sporulation frequencies in the experimentally evolved strains strains. The triangles represent control population, while squares represent fly treatment populations. The red fill is for the 2.5 day assay point, while the blue fill is for the 5 day assay point. Replicate populations in each treatment are modeled hierarchically.

**Figure 4:**
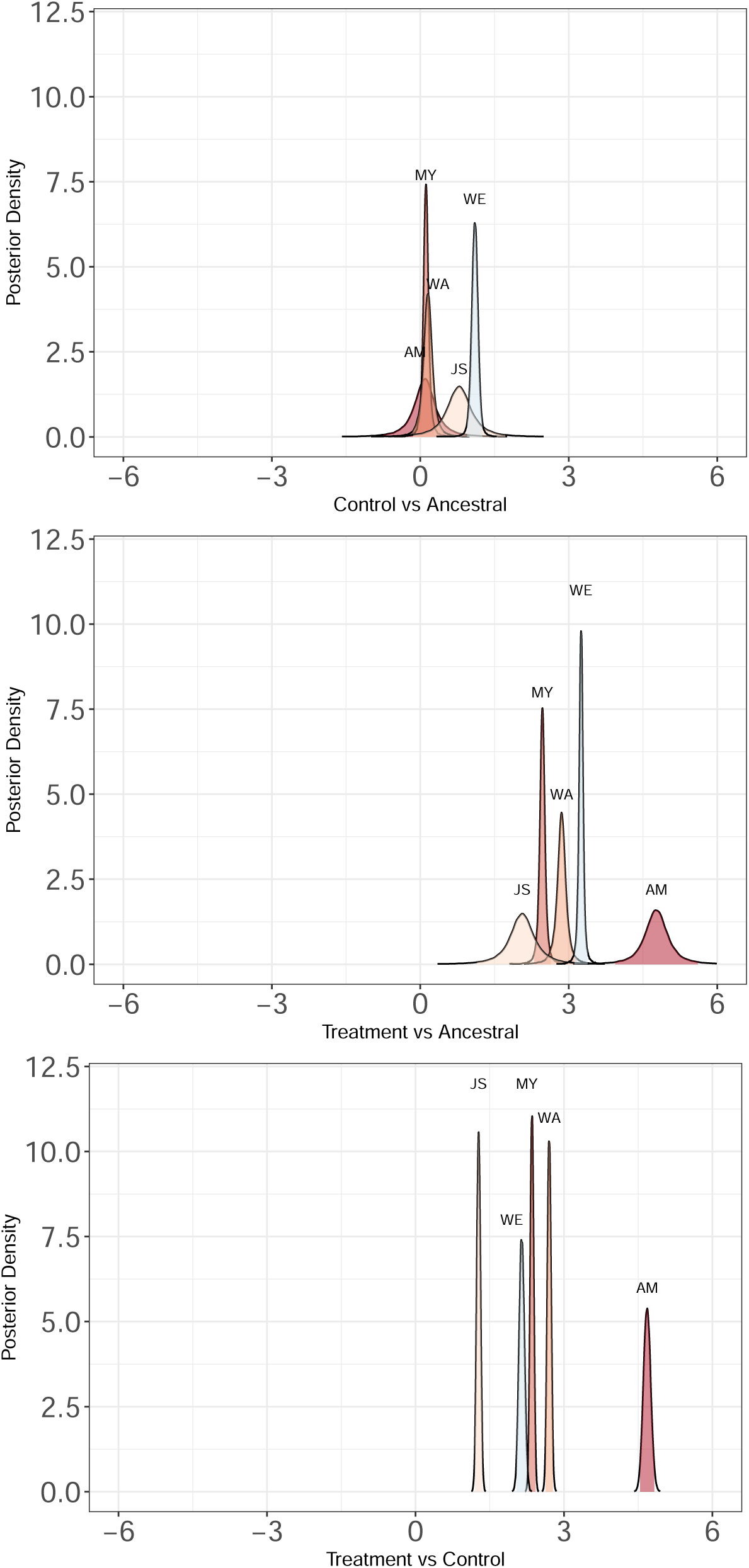
Three representations of the effect of treatment on sporulation rate. The top two panels compare the posterior distributions from the strain-specific models to the mean of the ancestral inference for that strain. The bottom panel compares the treatment to control for each strain.

The rate of sporulation completion, as measured by the assay time effect, varies with the degree of domestication (Figure 5). Wild strains showed a large response to fly vectoring in that there was a quicker evolved onset of sporulation (the posterior probability of the difference from the ancestral being less than zero is less than 10^−3^). The other three strains showed more modest responses to selection and the posterior probablity of a difference from the ancestor overlapped with 0.

**Figure 5:**
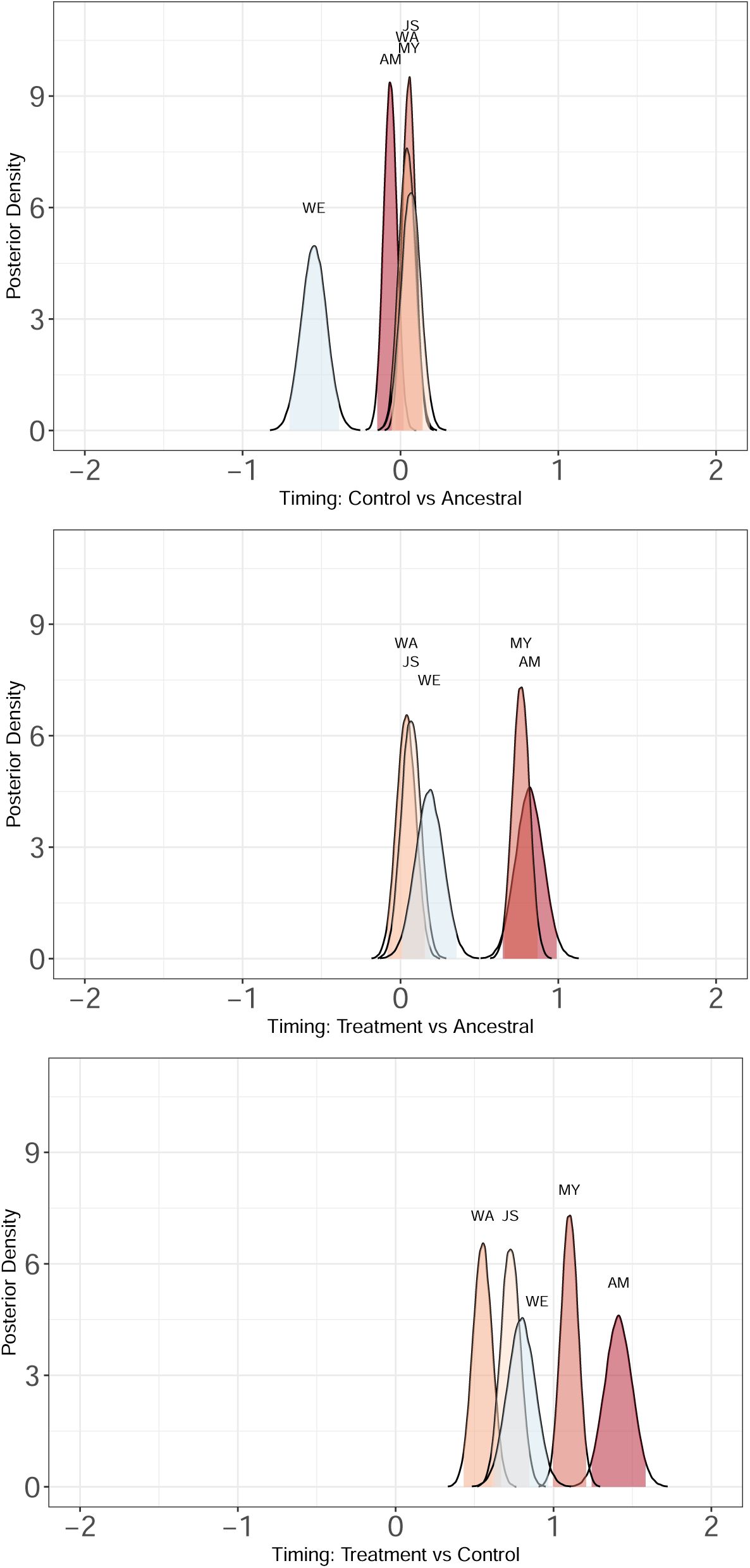
The effect of the fly treatment on the timing of sporulation. Larger positive values indicate sharper transition to sporulation between the time point at day 2.5 and at day 5. Comparisons between the treatment and control evolution experiments show that the transition to sporulation is sharper for populations evolving in the presence of flies.

### Selection responses in the control treatment

There is evidence that some of the control evolution treatment strains show a directional shift in sporulation rate (Figure 4). For the two domesticated strains (WE and JS) the posterior distributions for sporulation rates and assay time changed significantly after evolution. Both WE and JS increased their total sporulation rate in response to control evolution conditions (posterior probability of a change less than 0 of less than 10^−3^ for WE and of less than 0.05 for JS). In contrast, each of the other strains broadly overlapped with 0 change from the ancestor. In terms of the timing of sporulation, only WE was strongly diverged from the ancestor, with a decrease in the effect of assay time on sporulation (the posterior probability of the difference from the ancestral being greater than zero is less than 10^−3^).

### Comparison between the selection response in the fruit-fly and control conditions

Comparisons between the fly evolved treatment populations and the control evolved treatment populations show a consistent effect of exposure to fly vectoring that involved and increase in the total sporulation rate and an increase in the assay timing effect (see figures 4 and 5). In all cases the posterior probability of the difference between the treatment and control being less than zero is less than 10^−3^.

## Discussion

Previous studies have suggested that sporulation in budding yeast is an adaptation allowing lineages to survive passaging through insect vectors, e.g. (Coluccio *et al*., 2008; Neiman, 2005). These arguments are based on observations of differential survival by vegetative cells and spores in *Drosophila* frass. Other work has suggested that selection for dispersal traits, such as mating ability and germination, may also favor sporulation because insect digestion breaks up the ascus freeing non-related spores to mate following deposition of frass on fresh food sources (Reuter *et al*., 2007). Selection on sporulation onset and completion may however also depend on the timing of resource competition between unrelated vegetative cells, and the mortality effects of challenging environments (Ratcliff *et al*., 2013).

Our results mostly support the latest view about the adaptive value of sporulation. We found that strict passaging through the *Drosophila* digestive tract resulted in the evolution of both faster sporulation and higher sporulation rates. Since both of these traits are properties of the growing vegetative cells, we might have expected that domesticated strains would show partial loss of sensitivity to starvation as well as reduced initiation and completion of sporulation. Given that these strains are starting from a deficit in their tendency to sporulate, these lineages could have adapted to passaging through *Drosophila* guts by increasing vegetative cell survival through the gut. Indeed, analysis of a large set of yeast isolates has shown that some strains have evolved increased survival of quiescent vegetative cells (De Chiara *et al*., 2020). This was not the case here, where even the domesticated strains showed a strong response to the fly treatment in terms of their total sporulation rate. However, domesticated strains evolved a lower overall sporulation rate and lower speed of sporulation, as compared with the wild strains. This suggests that domestication led to the loss of mutational options that allow the cells to sense and respond to environmental changes. These results are reminiscent of those of Kvitek and Sherlock (Kvitek and Sherlock, 2013), where experimental evolution in constant environment led to the loss of developmental and physiological programs involved in the sensing of environmental variation.

Because our experiments allowed populations to grow for a fixed time period before ingestion, we expected selection for a steep change in sporulation rates associated with that timing or alternatively for vegetative cell starvation sensing. In particular, there was ample time before ingestion to deplete nutritional resources, sense it and respond appropriately. The results observed in the control populations, which were not passaged through the *Drosophila* gut but that were also subjected to selection for starvation resistance are instructive. In them, we found that only those derived from the domesticated isolates showed an increase in total sporulation rate. We might have expected all populations to evolve decreased sporulation if vegetative cell resistence instead evovled. Mutational options towards vegetative cell resistance thus appear to be fewer than those of sporulation. Selection responses in both of our treatments further indicate that the adaptive value at the origin of sporulation was not to survive insect vectoring, or outcrossing, but the ability to cope with challenging environments. Sporulation was perhaps just co-opted as a dispersal strategy because of ecological constraints, namely that small Drosophilids may preferentially diet on yeast (Schiabor *et al*., 2014) and are small enough to vector them between favorable habitats (Gibbs and Stanton, 2001; Stamps *et al*., 2012; Tsai *et al*., 2008).

We focused on selection for sporulation based on individual selection. Natural yeast populations are strongly spatially-structured and thus group-level selection must have also be at work in nature. Theoretical expectations for the evolution of sporulation in these circumstances depends on the way that cells gain resources from the environment and are passaged to future demes. In our experiments, there was strict vertical transmission of populations through insect ingestion and selection was strongest on surviving the passaging event itself. In spatially-heterogeneous conditions, sporulation by an individual cell should evolve in response to the pattern of variability in terms of both resource availability and timing of insect ingestion. Prior theory on this idea treats the evolution of the sporulation as a kin selection problem, asking how an individual cell that sporulates affects clone-mates in terms of resource availability (Ratcliff *et al*., 2013), because cells that sporulate will stop taking up resources, which are then available both to genetically-related and genetically-unrelated cells in the same environment. The degree of mixing between cell lineages in the founding of new demes probably limits kin selected benefits and will likely select for increases in the total number of viable emigrants each genotype produces from an existing demes; but such theory remains to be fully developed.

In the context of a competition-colonization trade-off (Tilman, 1994), a yeast strain that sporulates earlier or at a higher frequency before ingestion by insects is more likely to survive the process of ingestion, digestion and transfer. However, if the period of competition within a deme is long, the higher sporulating genotype will reproduce more slowly and eventually be displaced by genotypes that have lower sporulation rates. In contrast, once insect ingestion occurs, genotypes that have a higher fraction of cells in the sporulated state will have higher survivorship during vectoring and therefore increased representation in newly founded demes. Competition-colonization trade-offs can thus allow coexistence of alternative strategies, sometimes allowing many strategies to coexist (Snyder and Adler, 2011). We speculate that such competition-colonization trade-offs lead to coexistence of yeast species with similar physiological niches but differing sporulation and germination programs. In particular, *S. cerevisiae and* S. paradoxus have similar developmental and physiological profiles for surviving adverse environments but, tellingly, differing sporulation and germination programs (Murphy and Zeyl, 2010, 2012). A possible explanation for this coexistence therefore is that they occupy distinct positions in the competition-colonization space. And in general, such competition-colonization trade-offs may explain much of the biological diversity found in microbes.

## Supporting information

Supplemental Analysis

## Acknowledgements

We would like to acknowledge Fernanda Pett for her fantastic diagrams of the experimental design. We thank Uma Rajpurkar and Tracy Yu. HT is supported Agence Nationale pour la Recherche (ANR-17-CE02-0017-01, ANR-18-CE02-0017-01), SRP by the National Science Foundation (EF-1137835) and the pepiniere interdisciplinaire CNRS-PSL Eco-Evo-Devo.

