## Supplemental Analysis for "Testing the adaptive value of sporulation in budding yeast using experimental evolution"

*Stephen Proulx*

*February 23, 2020*

### Setup

Load the data file and recast the matrix as well as renameing some columns to make the names easier to parse.

```
spore_data <-read.csv("LTEE_DATA_FinalAssay.csv") %>%
  as_tibble() %>%
  select(-questions) %>%
  mutate(strain.number = as.integer(as.numeric(strain.region))) %>%
  mutate(strain.replicate.cat = as.factor(Strain.isolate.replicate)) %>%
  mutate(assay.number.cat = as.factor(Assay.Number)) %>%
  gather(time.point.cell.type,number.cells,number.sporulated.cells.at.2.5.days,
         number.sporulated.cells.at.5.days,number.vegetative.cells.at.2.5.days,
         number.vegetative.cells.at.5.days) %>%
  separate(time.point.cell.type,c("s1","s2","s3","s4","s5")) %>%
  mutate(cell.type = s2)%>%
  mutate(time.point = s5) %>%
  select(-s1,-s2,-s3,-s4,-s5) %>%
  spread(cell.type,number.cells) %>%
  mutate(assay.time = ifelse(time.point==2,2.5,5.0)) %>%
  rename(vegetative.count=vegetative) %>%
  rename(sporulated.count=sporulated) %>%
  select(-total.cells.counted.at.2.5.days,-percent.spores.at.2.5.days,
         -total.cells.counted.at.5.days,-percent.spores.at.5.days,
         -strain.replicate.cat,-time.point )  %>%
   arrange(treatment.type,strain.number,Ancestral.Population.ID) %>%
  mutate(strain.region = fct_relevel(strain.region,"AM","MY","WA","JS","WE"))
```

Load the pre-run BRMS models if they have already been saved.

```
load(file="StrainSpecificRandomEModels2.RData")
```

#### Exploratory plots

Make plots of the raw data for visualization.

```
mypalette<-brewer.pal(9,"Set1")
```

Plot all of the data by yeast strain and populaiton. The ancestral data are shown as a single box plot to make the visualization easier, and because the replicate population numbers in the ancestral treatment are not condordant with those of the two experimental treatments. The triangles are control experimental data and the squares are fly experimental data.

```
subdata<-spore_data %>% filter( assay.time==5 ) %>%
  mutate(replicate.pop.number =
           ((Ancestral.Population.ID==1)*1+(Ancestral.Population.ID==2)*2 +
  (Ancestral.Population.ID==11)*3 +(Ancestral.Population.ID==12)*4 +
    (Ancestral.Population.ID==3)*1+(Ancestral.Population.ID==4)*2 +
  (Ancestral.Population.ID==13)*3 +(Ancestral.Population.ID==14)*4+
  (Ancestral.Population.ID==5)*1+(Ancestral.Population.ID==6)*2 +
  (Ancestral.Population.ID==15)*3 +(Ancestral.Population.ID==16)*4+
  (Ancestral.Population.ID==7)*1+(Ancestral.Population.ID==8)*2 +
  (Ancestral.Population.ID==17)*3 +(Ancestral.Population.ID==18)*4+
  (Ancestral.Population.ID==9)*1+(Ancestral.Population.ID==10)*2 +
  (Ancestral.Population.ID==19)*3 +
    (Ancestral.Population.ID==20)*4)*(treatment.type!="ancestral"))%>%
  mutate(replicate.pop.number = replace_na(replicate.pop.number,0))

       ggplot(data = subdata,
       aes(x = strain.region,
           y = sporulated.count/(sporulated.count + vegetative.count),
       fill=as.factor(replicate.pop.number) )) +
         geom_boxplot(data=filter(subdata, treatment.type=="ancestral") ,
                      alpha=0.3, fill=mypalette[7], color="grey")     +
         geom_point(  data=filter(subdata, treatment.type=="control"),
                      aes(fill = mypalette[replicate.pop.number] ),
                      size = 3, shape = 24,
                      position = position_jitterdodge(jitter.width=.2,jitter.height=.001)) +
          geom_point(data=filter(subdata, treatment.type=="treatment"),
                     aes(fill = mypalette[replicate.pop.number]),
                     size = 3, shape = 22,
                     position = position_jitterdodge(jitter.width=.2,jitter.height=.001)) +
  scale_fill_brewer(palette="Spectral") +
           scale_y_continuous(limits = c(0,1) , breaks = c(0,.25,.5,.75,1.0))+
          labs(x = "Yeast Strain",
       y = "Fraction Spores" , fill="")+
  theme(text = element_text(size = 18),
        legend.position = "")
```

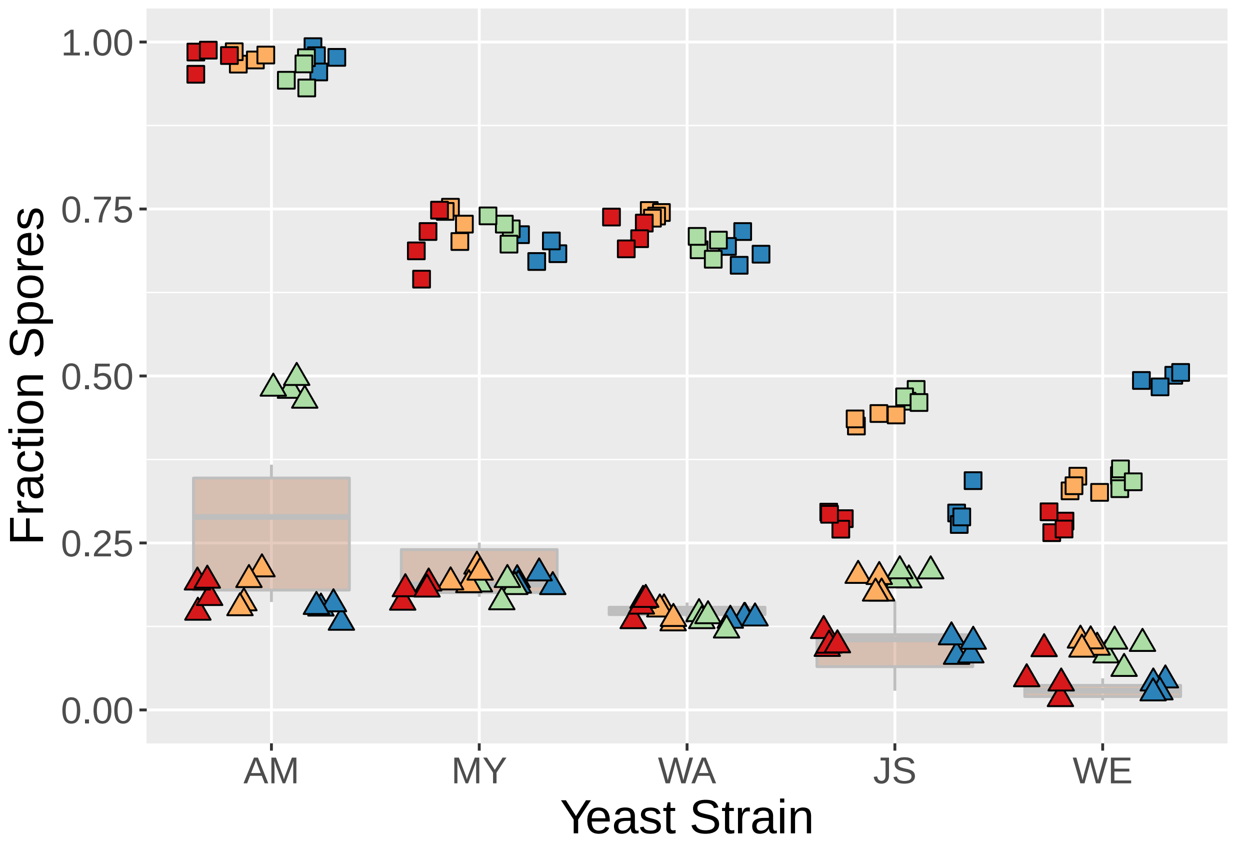


#### Bayesian fitting of the ancestral data

The ancestral data is fit with three alternative models. Becuase this is pre-run and the results are saved these are not evaluated.

Model AncNNull: Ancestral population replicate is nested within strain. Counts of sporulated cells (out of total cells counted) are binomially distrubted with a mean value that depends on the assay time, but not the strain. This is sort of a null model that we use to show that strain does have an effect.

```
mydata.sub<-spore_data %>%
    mutate(total = sporulated.count + vegetative.count) %>%
    as_tibble()  %>%
    mutate(strain.rep.id=    as.numeric(as.factor(Ancestral.Population.ID))) %>%
    mutate(strain.rep.id.0 = replace_na(strain.rep.id,0))%>%
    mutate(tratment.num = as.numeric(as.factor(treatment.type))) %>%
    mutate(assay.time.cat=as.factor(assay.time)) %>%
    mutate(techreps =    as.factor( Strain.isolate.replicate*10+technical.replicate) )%>%
    filter(treatment.type=="ancestral")

fitbin_brm_AncNNull<- brm(sporulated.count | trials(total) ~assay.time.cat +
                            assay.number.cat  + (strain.region|techreps) ,
                          data = mydata.sub,
                          family = binomial("logit"),future = FALSE  ,
                          chains=16 , iter = 6000 , seed=234124,
                          cores=16 ,control = list(adapt_delta = 0.99) )
```

Model AncNNoInteraction: Ancestral population replicate is nested within strain. Counts of sporulated cells (out of total cells counted) are binomially distrubted with a mean value that depends on the strain type and the assay time, but not their interaction.

```
mydata.sub<-spore_data %>%
    mutate(total = sporulated.count + vegetative.count) %>%
    as_tibble()  %>%
    mutate(strain.rep.id=    as.numeric(as.factor(Ancestral.Population.ID))) %>%
    mutate(strain.rep.id.0 = replace_na(strain.rep.id,0))%>%
    mutate(tratment.num = as.numeric(as.factor(treatment.type))) %>%
    mutate(assay.time.cat=as.factor(assay.time)) %>%
    mutate(techreps =    as.factor( Strain.isolate.replicate*10+technical.replicate) )%>%
    filter(treatment.type=="ancestral")

fitbin_brm_AncNNoInteraction<- brm(sporulated.count | trials(total) ~strain.region +
                                assay.time.cat + assay.number.cat  +
                                  (strain.region|techreps) ,
                                data = mydata.sub, family = binomial("logit"),
                                future = FALSE  ,chains=16 ,
                                iter = 6000 , seed=234124,
                                cores=16 ,control = list(adapt_delta = 0.99) )
```

Model AncN: Ancestral population replicate is nested within strain. Counts of sporulated cells (out of total cells counted) are binomially distrubted with a mean value that depends only on the strain type and its interaction with assay time.

```
mydata.sub<-spore_data %>%
    mutate(total = sporulated.count + vegetative.count) %>%
    as_tibble()  %>%
    mutate(strain.rep.id=    as.numeric(as.factor(Ancestral.Population.ID))) %>%
    mutate(strain.rep.id.0 = replace_na(strain.rep.id,0))%>%
    mutate(tratment.num = as.numeric(as.factor(treatment.type))) %>%
    mutate(assay.time.cat=as.factor(assay.time)) %>%
    mutate(techreps =    as.factor( Strain.isolate.replicate*10+technical.replicate) )%>%
    filter(treatment.type=="ancestral")

fitbin_brm_AncN<- brm(sporulated.count | trials(total) ~strain.region*assay.time.cat +
                       assay.number.cat  + (strain.region|techreps) ,
                     data = mydata.sub, family = binomial("logit"),
                     future = FALSE  ,chains=16 ,
                     iter = 12000 , seed=234124,
                     cores=16 ,control = list(adapt_delta = 0.99) )
```

Make a plot of the inferred sporulation rates for each strain and each time point.

```
#summary(fitbin_brm_AncN)


me <- marginal_effects(fitbin_brm_AncN ,effects = "strain.region:assay.time.cat" ,
                       re_formula = NA , points=TRUE)
```

```
## Setting the number of trials to 1 by default if not specified otherwise.
```

```
  plot(me, plot = FALSE)[[1]] +
    scale_x_discrete("Strain")+
    scale_y_continuous(limits=c(0,.3),"Frequency of Spores") +
    labs(colour = "Assay Time") +
```

```r
  theme(legend.position = "top")
```

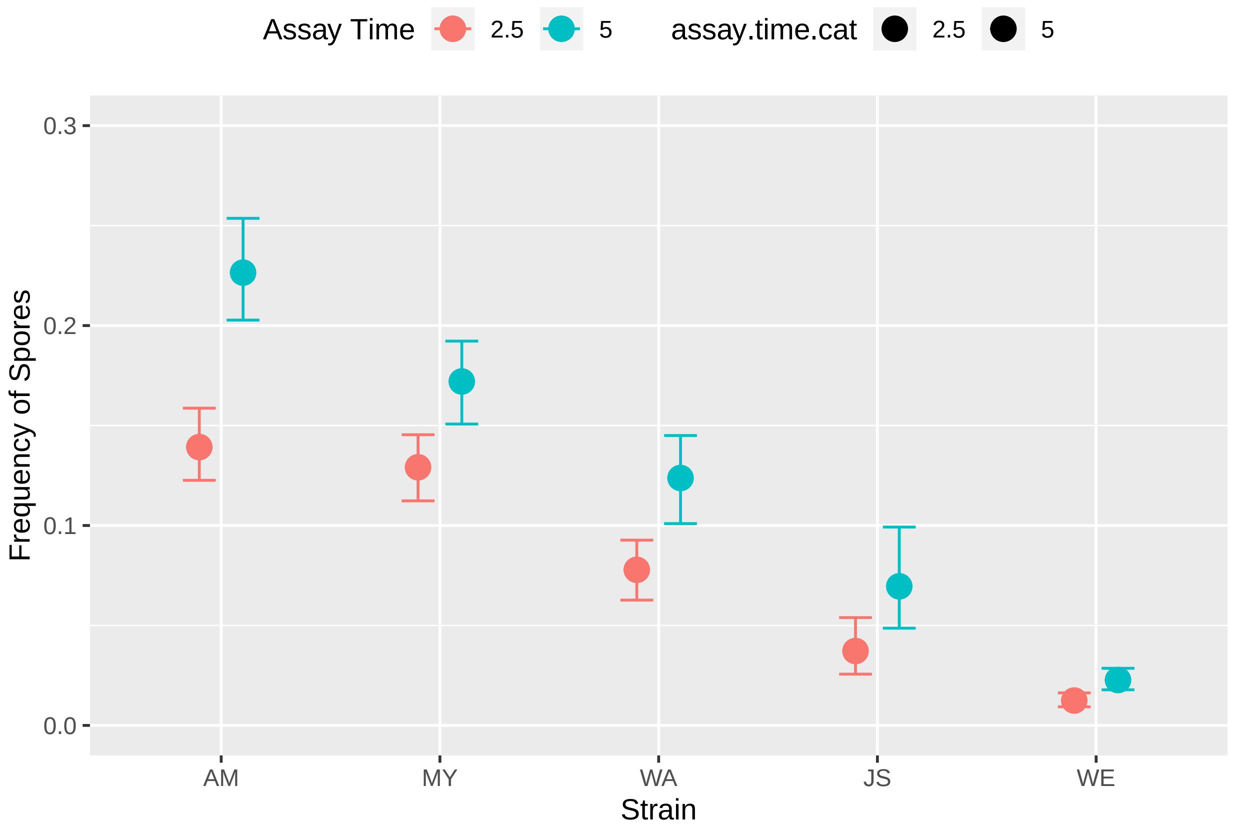


Test the alternative models for the ancestral populations. This shows that the model with an interaction between strain and timing of sporulation fits the data best.

```r
fitbin_brm_AncNNull<-add_criterion(fitbin_brm_AncNNull,c("loo","waic"))
fitbin_brm_AncNNoInteraction<-add_criterion(fitbin_brm_AncNNoInteraction,c("loo","waic"))
fitbin_brm_AncN<-add_criterion(fitbin_brm_AncN,c("loo","waic"))
```

```r
print(brms::loo_compare(fitbin_brm_AncNNull,fitbin_brm_AncNNoInteraction,
                        fitbin_brm_AncN, criterion = "loo") , simplify = FALSE)
```

```
##                              elpd_diff se_diff elpd_loo se_elpd_loo p_loo
## fitbin_brm_AncN                 0.0       0.0  -467.6     18.1        73.4
## fitbin_brm_AncNNoInteraction   -8.0       8.1  -475.5     18.2        73.7
## fitbin_brm_AncNNull           -15.6       8.4  -483.2     18.1        82.7
##                              se_p_loo looic  se_looic
## fitbin_brm_AncN                 9.7    935.2   36.2
## fitbin_brm_AncNNoInteraction    9.3    951.1   36.3
## fitbin_brm_AncNNull             9.6    966.3   36.1
```

```r
print(brms::loo_compare(fitbin_brm_AncNNull,fitbin_brm_AncNNoInteraction,
                        fitbin_brm_AncN, criterion = "waic") , simplify = FALSE)
```

```
##                              elpd_diff se_diff elpd_waic se_elpd_waic
## fitbin_brm_AncN                 0.0       0.0  -455.4      16.6
## fitbin_brm_AncNNoInteraction   -9.7       7.5  -465.2      16.9
## fitbin_brm_AncNNull           -13.9       7.8  -469.3      16.6
```

```
##                                p_waic se_p_waic waic   se_waic
## fitbin_brm_AncN                 61.3    8.0     910.9   33.3
## fitbin_brm_AncNNoInteraction   63.3    7.9     930.3   33.8
## fitbin_brm_AncNNull             68.8    7.9     938.6   33.1
```

Make posterior distribution plots for the difference between the strains in the ancestral model. This compares each strain with the strain that has the next lowest sporulation rate

```
samplesspec <-posterior_samples(fitbin_brm_AncN,"^b")%>%
  mutate(AM5=`b_assay.time.cat5`,
         MY5=`b_assay.time.cat5`+`b_strain.regionMY`+`b_strain.regionMY:assay.time.cat5`,
         WA5=`b_assay.time.cat5`+`b_strain.regionWA`+`b_strain.regionWA:assay.time.cat5`,
         JS5=`b_assay.time.cat5`+`b_strain.regionJS`+`b_strain.regionJS:assay.time.cat5`,
         WE5=`b_assay.time.cat5`+`b_strain.regionWE`+`b_strain.regionWE:assay.time.cat5`)



lq = 0.025
uq = 0.975
d1 <- density(samplesspec$AM5-samplesspec$MY5)
dd1 <- with(d1, data.frame(x, y)) %>% filter(y > 0.01)
qs1 = quantile(ecdf(samplesspec$AM5-samplesspec$MY5), prob = c(lq, 0.5, uq))
d2 <- density(samplesspec$MY5-samplesspec$WA5)
dd2 <- with(d2, data.frame(x, y)) %>% filter(y > 0.01)
qs2 = quantile(ecdf(samplesspec$MY5-samplesspec$WA5), prob = c(lq, 0.5, uq))
d3 <- density(samplesspec$WA5-samplesspec$JS5)
dd3 <- with(d3, data.frame(x, y)) %>% filter(y > 0.01)
qs3 = quantile(ecdf(samplesspec$WA5-samplesspec$JS5), prob = c(lq, 0.5, uq))
d4 <- density(samplesspec$JS5-samplesspec$WE5)
dd4 <- with(d4, data.frame(x, y)) %>% filter(y > 0.01)
qs4 = quantile(ecdf(samplesspec$JS5-samplesspec$WE5), prob = c(lq, 0.5, uq))





ggplot(data = dd1, aes(x, y)) +
  geom_line(data = dd1) + geom_ribbon(data = filter(dd1, x > qs1[[1]] & x < qs1[[3]]),
                                      aes(ymax = y), ymin = 0, fill = "#B2182B",
colour = NA, alpha = 0.5) +
  geom_line(data = dd2) + geom_ribbon(data = filter(dd2, x > qs2[[1]] & x < qs2[[3]]),
                                      aes(ymax = y), ymin = 0, fill = "#D6604D",
colour = NA, alpha = 0.5) +
  geom_line(data = dd3) + geom_ribbon(data = filter(dd3, x > qs3[[1]] & x < qs3[[3]]),
                                      aes(ymax = y), ymin = 0, fill = "#F4A582",
colour = NA, alpha = 0.5) +
  geom_line(data = dd4) + geom_ribbon(data = filter(dd4, x > qs4[[1]] & x < qs4[[3]]),
                                      aes(ymax = y), ymin = 0, fill = "#FDDBC7",
colour = NA, alpha = 0.5) +
    scale_y_continuous(limits = c(0,7), name = "Posterior Density") +
  scale_x_continuous(name ="Strain Comparisons",limits = c(-2,2) , breaks = c(-2,-1,0,1,2)) +
  annotate("text", x = c(qs1[[2]], qs2[[2]]+.2, qs3[[2]]+0.1, qs4[[2]]), y = c(6.7,5,2.3,2.2),
           label = c("AMvsMY","MYvsWA","WAvsJS","JSvsWE"), parse = TRUE, size = 3) +
  theme_bw() + theme(axis.text = element_text(family = "Helvetica",
size = 18), text = element_text(family = "Helvetica", size = 12))
```

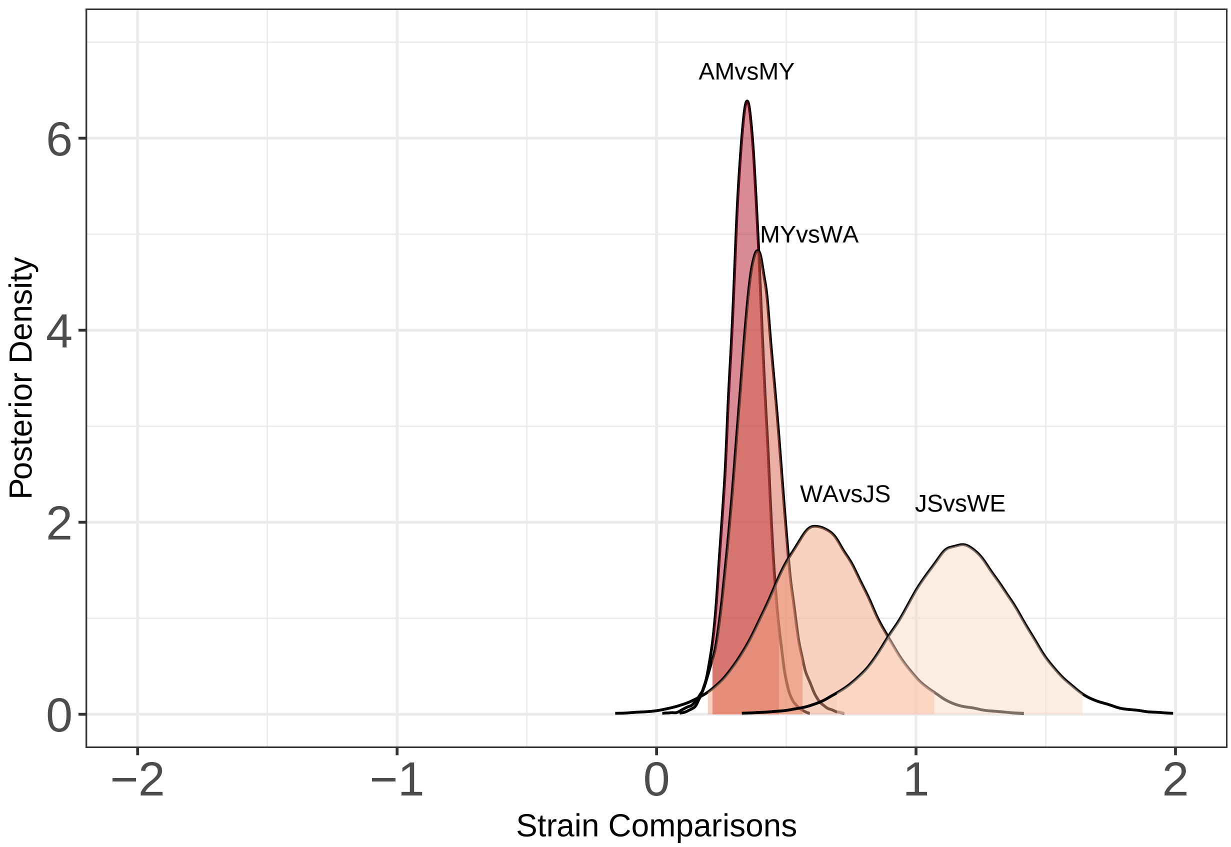


Determine the probability that the posterior distributions of the difference between strains overlaps with zero.

```
overlap <- tibble(comparison = c("AMvsMY","MYvsWA","WAvsJS","JSvsWE"),
                  prob = c(ecdf(samplesspec$AM5-samplesspec$MY5)(0),
                           ecdf(samplesspec$MY5-samplesspec$WA5)(0),
                           ecdf(samplesspec$WA5-samplesspec$JS5)(0),
                           ecdf(samplesspec$JS5-samplesspec$WE5)(0)  ))

print(overlap)
```

```
## # A tibble: 4 x 2
##   comparison      prob
##   <chr>          <dbl>
## 1 AMvsMY     0.0000208
## 2 MYvsWA     0.000167
## 3 WAvsJS     0.00438
## 4 JSvsWE     0.0000208
```

### Bayesian fitting for the experimental data

```
subdata<-spore_data  %>%
 mutate(replicate.pop.number =
          ((Ancestral.Population.ID==1)*1+(Ancestral.Population.ID==2)*2 +
  (Ancestral.Population.ID==11)*3 +(Ancestral.Population.ID==12)*4 +
    (Ancestral.Population.ID==3)*1+(Ancestral.Population.ID==4)*2 +
```

```
  (Ancestral.Population.ID==13)*3 +(Ancestral.Population.ID==14)*4+
  (Ancestral.Population.ID==5)*1+(Ancestral.Population.ID==6)*2 +
  (Ancestral.Population.ID==15)*3 +(Ancestral.Population.ID==16)*4+
  (Ancestral.Population.ID==7)*1+(Ancestral.Population.ID==8)*2 +
  (Ancestral.Population.ID==17)*3 +(Ancestral.Population.ID==18)*4+
  (Ancestral.Population.ID==9)*1+(Ancestral.Population.ID==10)*2 +
  (Ancestral.Population.ID==19)*3 +
    (Ancestral.Population.ID==20)*4)*(treatment.type!="ancestral"))%>%
  mutate(replicate.pop.number = replace_na(replicate.pop.number,0))


mydata.sub<-subdata %>%
    mutate(total = sporulated.count + vegetative.count) %>%
    as_tibble()  %>%
    mutate(strain.rep.id=    as.numeric(as.factor(Ancestral.Population.ID))) %>%
    mutate(strain.rep.id.0 = replace_na(strain.rep.id,0))%>%
    mutate(tratment.num = as.numeric(as.factor(treatment.type))) %>%
    mutate(assay.time.cat=as.factor(assay.time)) %>%
    mutate(Strain.isolate.replicate.cat = as.factor(Strain.isolate.replicate)) %>%
    mutate(techreps =    as.factor( replicate.pop.number*1) )%>%
    filter(treatment.type!="ancestral" , strain.region=="AM"  )




#random effect model
fitbin_brm_AMExpRandom<- brm(sporulated.count | trials(total) ~treatment.type*assay.time.cat+
                               (1|Strain.isolate.replicate.cat),
                                data = mydata.sub, family = binomial("logit"),
                             future = FALSE  ,chains=16 ,
                                iter = 12000 , seed=234124,
                             cores=16 ,control = list(adapt_delta = 0.99) )
```

```
 subdata<-spore_data  %>%
  mutate(replicate.pop.number =
           ((Ancestral.Population.ID==1)*1+(Ancestral.Population.ID==2)*2 +
  (Ancestral.Population.ID==11)*3 +(Ancestral.Population.ID==12)*4 +
    (Ancestral.Population.ID==3)*1+(Ancestral.Population.ID==4)*2 +
  (Ancestral.Population.ID==13)*3 +(Ancestral.Population.ID==14)*4+
  (Ancestral.Population.ID==5)*1+(Ancestral.Population.ID==6)*2 +
  (Ancestral.Population.ID==15)*3 +(Ancestral.Population.ID==16)*4+
  (Ancestral.Population.ID==7)*1+(Ancestral.Population.ID==8)*2 +
  (Ancestral.Population.ID==17)*3 +(Ancestral.Population.ID==18)*4+
  (Ancestral.Population.ID==9)*1+(Ancestral.Population.ID==10)*2 +
  (Ancestral.Population.ID==19)*3 +
    (Ancestral.Population.ID==20)*4)*(treatment.type!="ancestral"))%>%
  mutate(replicate.pop.number = replace_na(replicate.pop.number,0))


mydata.sub<-subdata %>%
    mutate(total = sporulated.count + vegetative.count) %>%
    as_tibble()  %>%
    mutate(strain.rep.id=    as.numeric(as.factor(Ancestral.Population.ID))) %>%
    mutate(strain.rep.id.0 = replace_na(strain.rep.id,0))%>%
```

```
    mutate(tratment.num = as.numeric(as.factor(treatment.type))) %>%
    mutate(assay.time.cat=as.factor(assay.time)) %>%
    mutate(Strain.isolate.replicate.cat = as.factor(Strain.isolate.replicate)) %>%
    mutate(techreps =    as.factor( replicate.pop.number*1) )%>%
    filter(treatment.type!="ancestral" , strain.region=="MY"  )




#random effect model
fitbin_brm_MYExpRandom<- brm(sporulated.count | trials(total) ~
                              treatment.type*assay.time.cat +
                              (1|Strain.isolate.replicate.cat),
                               data = mydata.sub, family = binomial("logit"),
                             future = FALSE  ,chains=16 ,
                               iter = 12000 , seed=234124, cores=16 ,
                             control = list(adapt_delta = 0.99) )
```

```
 subdata<-spore_data  %>%
  mutate(replicate.pop.number =
           ((Ancestral.Population.ID==1)*1+(Ancestral.Population.ID==2)*2 +
  (Ancestral.Population.ID==11)*3 +(Ancestral.Population.ID==12)*4 +
    (Ancestral.Population.ID==3)*1+(Ancestral.Population.ID==4)*2 +
  (Ancestral.Population.ID==13)*3 +(Ancestral.Population.ID==14)*4+
  (Ancestral.Population.ID==5)*1+(Ancestral.Population.ID==6)*2 +
  (Ancestral.Population.ID==15)*3 +(Ancestral.Population.ID==16)*4+
  (Ancestral.Population.ID==7)*1+(Ancestral.Population.ID==8)*2 +
  (Ancestral.Population.ID==17)*3 +(Ancestral.Population.ID==18)*4+
  (Ancestral.Population.ID==9)*1+(Ancestral.Population.ID==10)*2 +
  (Ancestral.Population.ID==19)*3 +
    (Ancestral.Population.ID==20)*4)*(treatment.type!="ancestral"))%>%
  mutate(replicate.pop.number = replace_na(replicate.pop.number,0))


mydata.sub<-subdata %>%
    mutate(total = sporulated.count + vegetative.count) %>%
    as_tibble()  %>%
    mutate(strain.rep.id=    as.numeric(as.factor(Ancestral.Population.ID))) %>%
    mutate(strain.rep.id.0 = replace_na(strain.rep.id,0))%>%
    mutate(tratment.num = as.numeric(as.factor(treatment.type))) %>%
    mutate(assay.time.cat=as.factor(assay.time)) %>%
    mutate(Strain.isolate.replicate.cat = as.factor(Strain.isolate.replicate)) %>%
    mutate(techreps =    as.factor( replicate.pop.number*1) )%>%
    filter(treatment.type!="ancestral" , strain.region=="WA"  )




#random effect model
fitbin_brm_WAExpRandom<- brm(sporulated.count | trials(total) ~
                              treatment.type*assay.time.cat +
                              (1|Strain.isolate.replicate.cat),
                               data = mydata.sub, family = binomial("logit"),
                             future = FALSE  ,chains=16 ,
```

```
                                iter = 12000 , seed=234124, cores=16 ,
                             control = list(adapt_delta = 0.99) )
```

```
 subdata<-spore_data  %>%
  mutate(replicate.pop.number =
           ((Ancestral.Population.ID==1)*1+(Ancestral.Population.ID==2)*2 +
  (Ancestral.Population.ID==11)*3 +(Ancestral.Population.ID==12)*4 +
    (Ancestral.Population.ID==3)*1+(Ancestral.Population.ID==4)*2 +
  (Ancestral.Population.ID==13)*3 +(Ancestral.Population.ID==14)*4+
  (Ancestral.Population.ID==5)*1+(Ancestral.Population.ID==6)*2 +
  (Ancestral.Population.ID==15)*3 +(Ancestral.Population.ID==16)*4+
  (Ancestral.Population.ID==7)*1+(Ancestral.Population.ID==8)*2 +
  (Ancestral.Population.ID==17)*3 +(Ancestral.Population.ID==18)*4+
  (Ancestral.Population.ID==9)*1+(Ancestral.Population.ID==10)*2 +
  (Ancestral.Population.ID==19)*3 +
    (Ancestral.Population.ID==20)*4)*(treatment.type!="ancestral"))%>%
  mutate(replicate.pop.number = replace_na(replicate.pop.number,0))


mydata.sub<-subdata %>%
    mutate(total = sporulated.count + vegetative.count) %>%
    as_tibble()  %>%
    mutate(strain.rep.id=    as.numeric(as.factor(Ancestral.Population.ID))) %>%
    mutate(strain.rep.id.0 = replace_na(strain.rep.id,0))%>%
    mutate(tratment.num = as.numeric(as.factor(treatment.type))) %>%
    mutate(assay.time.cat=as.factor(assay.time)) %>%
    mutate(Strain.isolate.replicate.cat = as.factor(Strain.isolate.replicate)) %>%
    mutate(techreps =    as.factor( replicate.pop.number*1) )%>%
    filter(treatment.type!="ancestral" , strain.region=="JS"  )




#random effect model
fitbin_brm_JSExpRandom<- brm(sporulated.count | trials(total) ~
                               treatment.type*assay.time.cat +
                               (1|Strain.isolate.replicate.cat),
                                data = mydata.sub, family = binomial("logit"),
                             future = FALSE  ,chains=16 ,
                                iter = 12000 , seed=234124, cores=16 ,
                             control = list(adapt_delta = 0.99) )
```

```
 subdata<-spore_data  %>%
  mutate(replicate.pop.number =
           ((Ancestral.Population.ID==1)*1+(Ancestral.Population.ID==2)*2 +
  (Ancestral.Population.ID==11)*3 +(Ancestral.Population.ID==12)*4 +
    (Ancestral.Population.ID==3)*1+(Ancestral.Population.ID==4)*2 +
  (Ancestral.Population.ID==13)*3 +(Ancestral.Population.ID==14)*4+
  (Ancestral.Population.ID==5)*1+(Ancestral.Population.ID==6)*2 +
  (Ancestral.Population.ID==15)*3 +(Ancestral.Population.ID==16)*4+
  (Ancestral.Population.ID==7)*1+(Ancestral.Population.ID==8)*2 +
  (Ancestral.Population.ID==17)*3 +(Ancestral.Population.ID==18)*4+
  (Ancestral.Population.ID==9)*1+(Ancestral.Population.ID==10)*2 +
  (Ancestral.Population.ID==19)*3 +
```

```
   (Ancestral.Population.ID==20)*4)*(treatment.type!="ancestral"))%>%
  mutate(replicate.pop.number = replace_na(replicate.pop.number,0))


mydata.sub<-subdata %>%
    mutate(total = sporulated.count + vegetative.count) %>%
    as_tibble()  %>%
    mutate(strain.rep.id=    as.numeric(as.factor(Ancestral.Population.ID))) %>%
    mutate(strain.rep.id.0 = replace_na(strain.rep.id,0))%>%
    mutate(tratment.num = as.numeric(as.factor(treatment.type))) %>%
    mutate(assay.time.cat=as.factor(assay.time)) %>%
    mutate(Strain.isolate.replicate.cat = as.factor(Strain.isolate.replicate)) %>%
    mutate(techreps =    as.factor( replicate.pop.number*1) )%>%
    filter(treatment.type!="ancestral" , strain.region=="WE"  )




#random effect model
fitbin_brm_WEExpRandom<- brm(sporulated.count | trials(total) ~
                               treatment.type*assay.time.cat +
                               (1|Strain.isolate.replicate.cat),
                                data = mydata.sub, family = binomial("logit"),
                             future = FALSE  ,chains=16 ,
                                iter = 12000 , seed=234124, cores=16 ,
                             control = list(adapt_delta = 0.99) )
```

#### Plots and comparisons of the experimental data

```
cond<-tibble(treatment.type="treatment")



me <- marginal_effects(fitbin_brm_AMExpRandom ,effects = "treatment.type:assay.time.cat"  ,
                       conditions=cond, re_formula = NA , points=TRUE)
```

```
## Setting the number of trials to 1 by default if not specified otherwise.
```

```
conditionaldataAM=tibble(strain.region="AM",
    treatment.type=c("control","control","treatment","treatment"),
    assay.time.cat=c("2.5","5","2.5","5"),
    estimates = me$`treatment.type:assay.time.cat`$estimate__,
    lowers= me$`treatment.type:assay.time.cat`$lower__,
    uppers= me$`treatment.type:assay.time.cat`$upper__ )


me <- marginal_effects(fitbin_brm_MYExpRandom ,
                       effects = "treatment.type:assay.time.cat"  ,
                       conditions=cond, re_formula = NA , points=TRUE)
```

```
## Setting the number of trials to 1 by default if not specified otherwise.
```

```
conditionaldataMY=tibble(strain.region="MY",
    treatment.type=c("control","control","treatment","treatment"),
    assay.time.cat=c("2.5","5","2.5","5"),
    estimates = me$`treatment.type:assay.time.cat`$estimate__,
    lowers= me$`treatment.type:assay.time.cat`$lower__,
    uppers= me$`treatment.type:assay.time.cat`$upper__ )


me <- marginal_effects(fitbin_brm_WAExpRandom ,
                       effects = "treatment.type:assay.time.cat"  ,
                       conditions=cond, re_formula = NA , points=TRUE)
```

```
## Setting the number of trials to 1 by default if not specified otherwise.
```

```
conditionaldataWA=tibble(strain.region="WA",
  treatment.type=c("control","control","treatment","treatment"),
  assay.time.cat=c("2.5","5","2.5","5"),
  estimates = me$`treatment.type:assay.time.cat`$estimate__,
  lowers= me$`treatment.type:assay.time.cat`$lower__,
  uppers= me$`treatment.type:assay.time.cat`$upper__ )


me <- marginal_effects(fitbin_brm_JSExpRandom ,
                       effects = "treatment.type:assay.time.cat"  ,
                       conditions=cond, re_formula = NA , points=TRUE)
```

```
## Setting the number of trials to 1 by default if not specified otherwise.
```

```
conditionaldataJS=tibble(strain.region="JS",
    treatment.type=c("control","control","treatment","treatment"),
    assay.time.cat=c("2.5","5","2.5","5"),
    estimates = me$`treatment.type:assay.time.cat`$estimate__,
    lowers= me$`treatment.type:assay.time.cat`$lower__,
    uppers= me$`treatment.type:assay.time.cat`$upper__ )


me <- marginal_effects(fitbin_brm_WEExpRandom ,
                       effects = "treatment.type:assay.time.cat"  ,
                       conditions=cond, re_formula = NA , points=TRUE)
```

```
## Setting the number of trials to 1 by default if not specified otherwise.
```

```
conditionaldataWE=tibble(strain.region="WE",
    treatment.type=c("control","control","treatment","treatment"),
    assay.time.cat=c("2.5","5","2.5","5"),
    estimates = me$`treatment.type:assay.time.cat`$estimate__,
    lowers= me$`treatment.type:assay.time.cat`$lower__,
    uppers= me$`treatment.type:assay.time.cat`$upper__ )


conditionaldata <- rbind(conditionaldataAM,conditionaldataMY,
                         conditionaldataWA,conditionaldataJS,conditionaldataWE) %>%
  mutate(strain.region = fct_relevel(strain.region,"AM","MY","WA","JS","WE"))
```

```r
ggplot(data=conditionaldata,
       aes(x=strain.region, y=estimates, color=assay.time.cat, fill=assay.time.cat)) +
  geom_errorbar(data=filter(conditionaldata,treatment.type=="control",assay.time.cat=="2.5"),
                aes(ymin=lowers, ymax=uppers) ,  position= position_nudge(x=-0.25)  , width=0.1 )+
  geom_point(data=filter(conditionaldata,treatment.type=="control",assay.time.cat=="2.5"),
             position = position_nudge(x=-0.25) ,size = 3, shape = 24)+
  geom_errorbar(data=filter(conditionaldata,treatment.type=="control",assay.time.cat=="5"),
                aes(ymin=lowers, ymax=uppers) ,  position= position_nudge(x=-0.125)  , width=0.1 ) +
  geom_point(data=filter(conditionaldata,treatment.type=="control",assay.time.cat=="5"),
             position = position_nudge(x=-0.125) ,size = 3, shape = 24)+
  geom_errorbar(data=filter(conditionaldata,treatment.type=="treatment",assay.time.cat=="2.5"),
                aes(ymin=lowers, ymax=uppers) ,  position= position_nudge(x=0.125)  , width=0.1 )+
  geom_point(data=filter(conditionaldata,treatment.type=="treatment",assay.time.cat=="2.5"),
             position = position_nudge(x=0.125) ,size = 3, shape = 22)+
  geom_errorbar(data=filter(conditionaldata,treatment.type=="treatment",assay.time.cat=="5"),
                aes(ymin=lowers, ymax=uppers) ,  position= position_nudge(x=0.25)  , width=0.1 ) +
  geom_point(data=filter(conditionaldata,treatment.type=="treatment",assay.time.cat=="5"),
             position = position_nudge(x=0.25) ,size = 3, shape = 22)+
           scale_y_continuous(limits = c(0,1) , breaks = c(0,.25,.5,.75,1.0))+
          labs(x = "Yeast Strain",
      y = "Fraction Spores" , fill="")+
  theme(text = element_text(size = 18),
        legend.position = "")
```

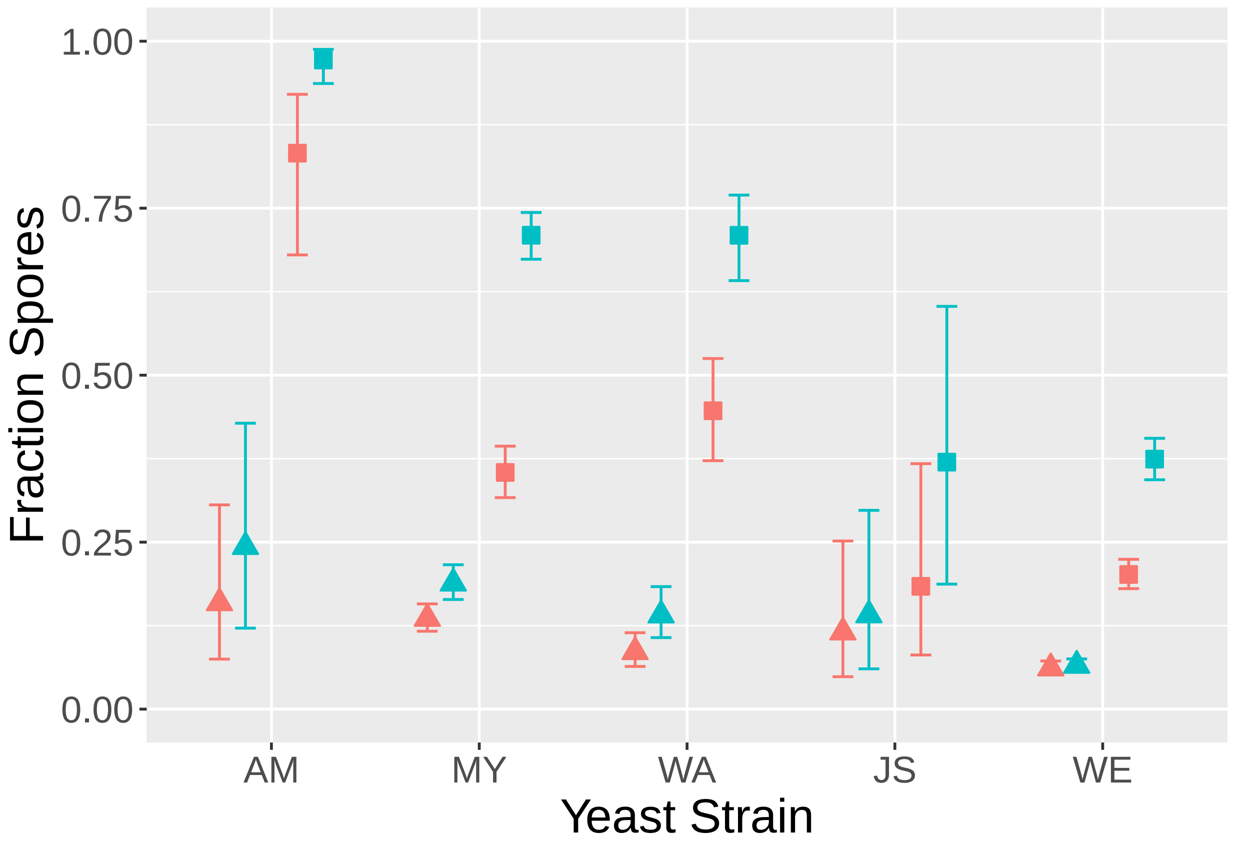

Extract the appropriate posterior distribution parmaeters for later comparison.

```
samplesAncestral <-posterior_samples(fitbin_brm_AncN,"^b")%>%
  mutate(AMAnc=`b_Intercept`+`b_assay.time.cat5`,
         MYAnc=`b_Intercept`+`b_strain.regionMY`+
           `b_assay.time.cat5`+`b_strain.regionMY:assay.time.cat5`,
         WAAnc=`b_Intercept`+`b_strain.regionWA`+
           `b_assay.time.cat5`+`b_strain.regionWA:assay.time.cat5`,
         JSAnc=`b_Intercept`+`b_strain.regionJS`+
           `b_assay.time.cat5`+`b_strain.regionJS:assay.time.cat5`,
         WEAnc=`b_Intercept`+`b_strain.regionWE`+
           `b_assay.time.cat5`+`b_strain.regionWE:assay.time.cat5`,
         AMAncTime=`b_assay.time.cat5`,
         MYAncTime=`b_assay.time.cat5`+`b_strain.regionMY:assay.time.cat5`,
         WAAncTime=`b_assay.time.cat5`+`b_strain.regionWA:assay.time.cat5`,
         JSAncTime=`b_assay.time.cat5`+`b_strain.regionJS:assay.time.cat5`,
         WEAncTime=`b_assay.time.cat5`+`b_strain.regionWE:assay.time.cat5`) %>%
  select(AMAnc,MYAnc,WAAnc,JSAnc,WEAnc,AMAncTime,MYAncTime,WAAncTime,JSAncTime,WEAncTime)

samplesAM <-posterior_samples(fitbin_brm_AMExpRandom,"^b")%>%
  mutate(AMControl2.5=`b_Intercept`,
         AMControl5=`b_Intercept`+`b_assay.time.cat5`,
         AMTreatment2.5=`b_Intercept`+`b_treatment.typetreatment`,
         AMTreatment5=`b_Intercept`+`b_treatment.typetreatment`+
           `b_assay.time.cat5`+`b_treatment.typetreatment:assay.time.cat5`,
         AMTTiming=`b_treatment.typetreatment:assay.time.cat5`,
         AMCTiming=`b_assay.time.cat5`)%>% as_tibble()%>%
  select(AMControl2.5,AMControl5,AMTreatment2.5,AMTreatment5,AMTTiming,AMCTiming)


samplesMY <-posterior_samples(fitbin_brm_MYExpRandom,"^b")%>%
  mutate(MYControl2.5=`b_Intercept` ,
         MYControl5=`b_Intercept`+ `b_assay.time.cat5`,
         MYTreatment2.5=`b_Intercept`+`b_treatment.typetreatment`,
         MYTreatment5=`b_Intercept`+`b_treatment.typetreatment`+
           `b_assay.time.cat5`+`b_treatment.typetreatment:assay.time.cat5`,
         MYTTiming=`b_treatment.typetreatment:assay.time.cat5`,
         MYCTiming=`b_assay.time.cat5`)%>%as_tibble()%>%
  select(MYControl2.5,MYControl5,MYTreatment2.5,MYTreatment5,MYTTiming,MYCTiming)


samplesWA <-posterior_samples(fitbin_brm_WAExpRandom,"^b")%>%
  mutate(WAControl2.5=`b_Intercept`,
         WAControl5=`b_Intercept`+`b_assay.time.cat5`,
         WATreatment2.5=`b_Intercept`+`b_treatment.typetreatment`,
         WATreatment5=`b_Intercept`+`b_treatment.typetreatment`+
           `b_assay.time.cat5`+`b_treatment.typetreatment:assay.time.cat5`,
         WATTiming=`b_treatment.typetreatment:assay.time.cat5`,
         WACTiming=`b_assay.time.cat5`)%>%as_tibble()%>%
  select(WAControl2.5,WAControl5,WATreatment2.5,WATreatment5,WATTiming,WACTiming)

samplesJS <-posterior_samples(fitbin_brm_JSExpRandom,"^b")%>%
  mutate(JSControl2.5=`b_Intercept`,
         JSControl5=`b_Intercept`+`b_assay.time.cat5`,
```

```
         JSTreatment2.5=`b_Intercept`+`b_treatment.typetreatment`,
         JSTreatment5=`b_Intercept`+`b_treatment.typetreatment`+
           `b_assay.time.cat5`+`b_treatment.typetreatment:assay.time.cat5`,
         JSTTiming=`b_treatment.typetreatment:assay.time.cat5`,
         JSCTiming=`b_treatment.typetreatment:assay.time.cat5`)%>%as_tibble()%>%
  select(JSControl2.5,JSControl5,JSTreatment2.5,JSTreatment5,JSTTiming,JSCTiming)

samplesWE <-posterior_samples(fitbin_brm_WEExpRandom,"^b")%>%
  mutate(WEControl2.5=`b_Intercept`,
         WEControl5=`b_Intercept`+`b_assay.time.cat5`,
         WETreatment2.5=`b_Intercept`+`b_treatment.typetreatment`,
         WETreatment5=`b_Intercept`+`b_treatment.typetreatment`+
           `b_assay.time.cat5`+`b_treatment.typetreatment:assay.time.cat5`,
         WETTiming=`b_treatment.typetreatment:assay.time.cat5`,
         WECTiming=`b_assay.time.cat5`)%>%as_tibble()%>%
  select(WEControl2.5,WEControl5,WETreatment2.5,WETreatment5,WETTiming,WECTiming)
```

```
#use the mean of the ancestral and the full posterior of the experimental
lq = 0.025
uq = 0.975
del1 <- (mean(samplesAncestral$AMAnc) - samplesAM$AMControl5)*(-1)
d1 <- density(del1)
dd1 <- with(d1, data.frame(x, y)) %>% filter(y > 0.01)
qs1 = quantile(ecdf( del1), prob = c(lq, 0.5, uq))
del2 <- (mean(samplesAncestral$MYAnc) - samplesMY$MYControl5)*(-1)
d2 <- density( del2)
dd2 <- with(d2, data.frame(x, y)) %>% filter(y > 0.01)
qs2 = quantile(ecdf(del2), prob = c(lq, 0.5, uq))
del3 <- (mean(samplesAncestral$WAAnc) - samplesWA$WAControl5)*(-1)
d3 <- density( del3)
dd3 <- with(d3, data.frame(x, y)) %>% filter(y > 0.01)
qs3 = quantile(ecdf(del3), prob = c(lq, 0.5, uq))
del4 <- (mean(samplesAncestral$JSAnc) - samplesJS$JSControl5)*(-1)
d4 <- density( del4)
dd4 <- with(d4, data.frame(x, y)) %>% filter(y > 0.01)
qs4 = quantile(ecdf(del4), prob = c(lq, 0.5, uq))
del5 <- (mean(samplesAncestral$WEAnc) - samplesWE$WEControl5)*(-1)
d5 <- density( del5)
dd5 <- with(d5, data.frame(x, y)) %>% filter(y > 0.01)
qs5 = quantile(ecdf(del5), prob = c(lq, 0.5, uq))






ggplot(data = dd1, aes(x, y)) +
  geom_line(data = dd1) + geom_ribbon(data = filter(dd1, x > qs1[[1]] & x < qs1[[3]]),
                                      aes(ymax = y), ymin = 0, fill = "#B2182B",
colour = NA, alpha = 0.5) +
  geom_line(data = dd2) + geom_ribbon(data = filter(dd2, x > qs2[[1]] & x < qs2[[3]]),
                                      aes(ymax = y), ymin = 0, fill = "#D6604D",
colour = NA, alpha = 0.5) +
    geom_line(data = dd3) + geom_ribbon(data = filter(dd3, x > qs3[[1]] & x < qs3[[3]]),
```

```
                                        aes(ymax = y), ymin = 0, fill = "#F4A582",
colour = NA, alpha = 0.5) +
  geom_line(data = dd4) + geom_ribbon(data = filter(dd4, x > qs4[[1]] & x < qs4[[3]]),
                                      aes(ymax = y), ymin = 0, fill = "#FDDBC7",
colour = NA, alpha = 0.5) +
  geom_line(data = dd5) + geom_ribbon(data = filter(dd5, x > qs5[[1]] & x < qs5[[3]]),
                                      aes(ymax = y), ymin = 0, fill = "#D1E5F0",
colour = NA, alpha = 0.5) +
    scale_y_continuous(limits = c(0,12), name = "Posterior Density") +
  scale_x_continuous(name ="Control vs Ancestral",limits = c(-6,6) ) +
  annotate("text", x = c(qs1[[2]]-.2, qs2[[2]], qs3[[2]]+.2, qs4[[2]], qs5[[2]]),
           y = c(2.5,7.7,4.5,2,7), label = c("AM","MY","WA","JS","WE"),
           parse = TRUE, size = 3) + theme_bw() + theme(axis.text = element_text(family = "Helvetica",
size = 18), text = element_text(family = "Helvetica", size = 12))
```

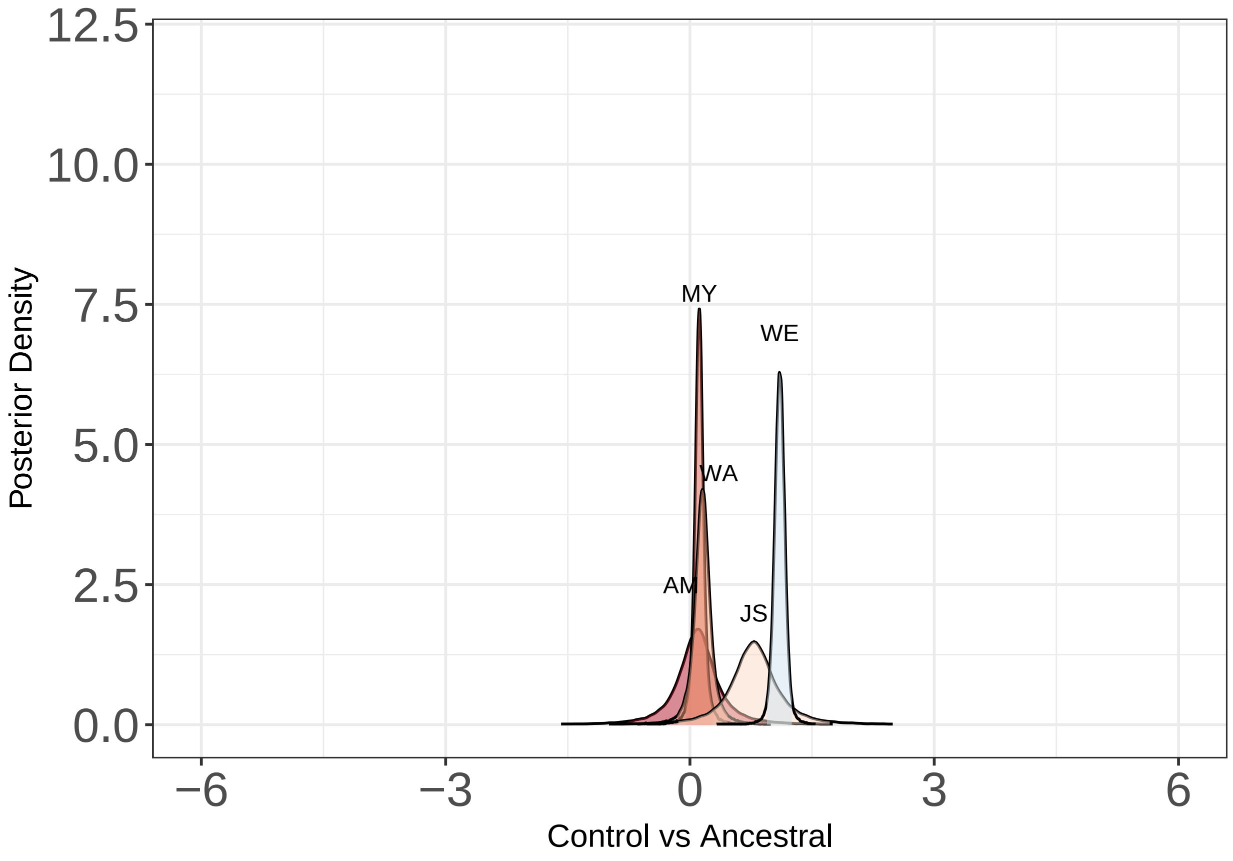


```
overlap <- tibble(strain = c("AM","MY","WA","JS","WE"),
                  prob = c( ecdf( del1)(0),ecdf( del2)(0),
                        ecdf( del3)(0),ecdf( del4)(0),ecdf( del5)(0) ))

print(overlap)
```

```
## # A tibble: 5 x 2
##   strain   prob
##   <chr>   <dbl>
## 1 AM     0.350
## 2 MY     0.0542
```

```
## 3 WA     0.0942
## 4 JS     0.0363
## 5 WE     0
```

```
lq = 0.025
uq = 0.975
del1 <- (mean(samplesAncestral$AMAnc) - samplesAM$AMTreatment5)*(-1)
d1 <- density(del1)
dd1 <- with(d1, data.frame(x, y)) %>% filter(y > 0.01)
qs1 = quantile(ecdf( del1), prob = c(lq, 0.5, uq))
del2 <- (mean(samplesAncestral$MYAnc) - samplesMY$MYTreatment5)*(-1)
d2 <- density( del2)
dd2 <- with(d2, data.frame(x, y)) %>% filter(y > 0.01)
qs2 = quantile(ecdf(del2), prob = c(lq, 0.5, uq))
del3 <- (mean(samplesAncestral$WAAnc) - samplesWA$WATreatment5)*(-1)
d3 <- density( del3)
dd3 <- with(d3, data.frame(x, y)) %>% filter(y > 0.01)
qs3 = quantile(ecdf(del3), prob = c(lq, 0.5, uq))
del4 <- (mean(samplesAncestral$JSAnc) - samplesJS$JSTreatment5)*(-1)
d4 <- density( del4)
dd4 <- with(d4, data.frame(x, y)) %>% filter(y > 0.01)
qs4 = quantile(ecdf(del4), prob = c(lq, 0.5, uq))
del5 <- (mean(samplesAncestral$WEAnc) - samplesWE$WETreatment5)*(-1)
d5 <- density( del5)
dd5 <- with(d5, data.frame(x, y)) %>% filter(y > 0.01)
qs5 = quantile(ecdf(del5), prob = c(lq, 0.5, uq))





ggplot(data = dd1, aes(x, y)) +
  geom_line(data = dd1) + geom_ribbon(data = filter(dd1, x > qs1[[1]] & x < qs1[[3]]),
                                      aes(ymax = y), ymin = 0, fill = "#B2182B",
colour = NA, alpha = 0.5) +
  geom_line(data = dd2) + geom_ribbon(data = filter(dd2, x > qs2[[1]] & x < qs2[[3]]),
                                      aes(ymax = y), ymin = 0, fill = "#D6604D",
colour = NA, alpha = 0.5) +
    geom_line(data = dd3) + geom_ribbon(data = filter(dd3, x > qs3[[1]] & x < qs3[[3]]),
                                        aes(ymax = y), ymin = 0, fill = "#F4A582",
colour = NA, alpha = 0.5) +
  geom_line(data = dd4) + geom_ribbon(data = filter(dd4, x > qs4[[1]] & x < qs4[[3]]),
                                      aes(ymax = y), ymin = 0, fill = "#FDDBC7",
colour = NA, alpha = 0.5) +
  geom_line(data = dd5) + geom_ribbon(data = filter(dd5, x > qs5[[1]] & x < qs5[[3]]),
                                      aes(ymax = y), ymin = 0, fill = "#D1E5F0",
colour = NA, alpha = 0.5) +
    scale_y_continuous(limits = c(0,12), name = "Posterior Density") +
  scale_x_continuous(name ="Treatment vs Ancestral",limits = c(-6,6) ) +
  annotate("text", x = c(qs1[[2]], qs2[[2]], qs3[[2]], qs4[[2]], qs5[[2]]),
           y = c(2.2,8,5,2.2,11), label = c("AM","MY","WA","JS","WE"),
           parse = TRUE, size = 3) + theme_bw() + theme(axis.text = element_text(family = "Helvetica",
size = 18), text = element_text(family = "Helvetica", size = 12))
```

```
## Warning: Removed 24 rows containing missing values (geom_path).
```

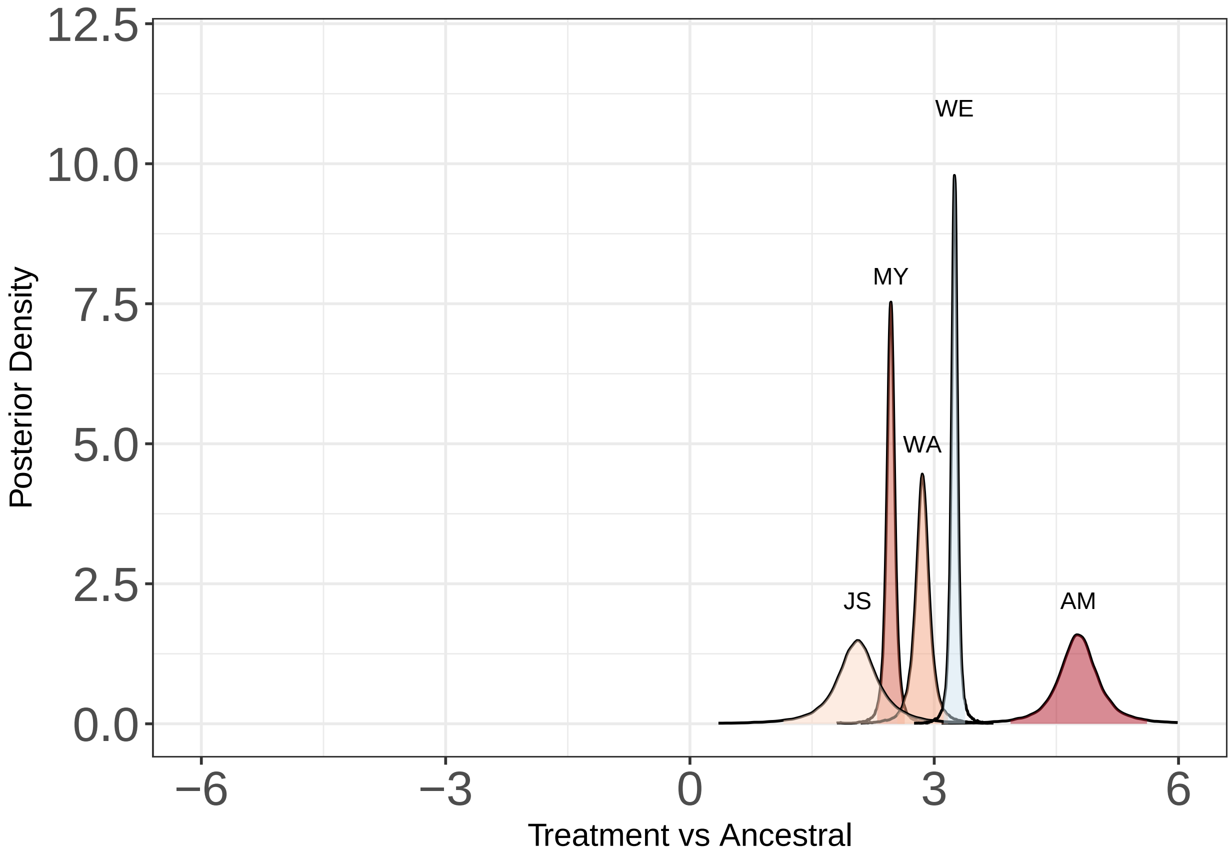


```
overlap <- tibble(strain = c("AM","MY","WA","JS","WE"),
                  prob = c( ecdf( del1)(0),ecdf( del2)(0),
                        ecdf( del3)(0),ecdf( del4)(0),ecdf( del5)(0) ))

print(overlap)
```

```
## # A tibble: 5 x 2
##   strain      prob
##   <chr>      <dbl>
## 1 AM     0
## 2 MY     0
## 3 WA     0.0000417
## 4 JS     0.00323
## 5 WE     0
```

```
lq = 0.025
uq = 0.975
del1 <- (samplesAM$AMControl5 - samplesAM$AMTreatment5)*(-1)
d1 <- density(del1)
dd1 <- with(d1, data.frame(x, y)) %>% filter(y > 0.01)
qs1 = quantile(ecdf( del1), prob = c(lq, 0.5, uq))
del2 <- (samplesMY$MYControl5 - samplesMY$MYTreatment5)*(-1)
d2 <- density( del2)
dd2 <- with(d2, data.frame(x, y)) %>% filter(y > 0.01)
qs2 = quantile(ecdf(del2), prob = c(lq, 0.5, uq))
del3 <- ( samplesWA$WAControl5 - samplesWA$WATreatment5)*(-1)
d3 <- density( del3)
```

```
dd3 <- with(d3, data.frame(x, y)) %>% filter(y > 0.01)
qs3 = quantile(ecdf(del3), prob = c(lq, 0.5, uq))
del4 <- ( samplesJS$JSControl5 - samplesJS$JSTreatment5)*(-1)
d4 <- density( del4)
dd4 <- with(d4, data.frame(x, y)) %>% filter(y > 0.01)
qs4 = quantile(ecdf(del4), prob = c(lq, 0.5, uq))
del5 <- (samplesWE$WEControl5 - samplesWE$WETreatment5)*(-1)
d5 <- density( del5)
dd5 <- with(d5, data.frame(x, y)) %>% filter(y > 0.01)
qs5 = quantile(ecdf(del5), prob = c(lq, 0.5, uq))





ggplot(data = dd1, aes(x, y)) +
  geom_line(data = dd1) + geom_ribbon(data = filter(dd1, x > qs1[[1]] & x < qs1[[3]]),
                                      aes(ymax = y), ymin = 0, fill = "#B2182B",
colour = NA, alpha = 0.5) +
  geom_line(data = dd2) + geom_ribbon(data = filter(dd2, x > qs2[[1]] & x < qs2[[3]]),
                                      aes(ymax = y), ymin = 0, fill = "#D6604D",
colour = NA, alpha = 0.5) +
    geom_line(data = dd3) + geom_ribbon(data = filter(dd3, x > qs3[[1]] & x < qs3[[3]]),
                                        aes(ymax = y), ymin = 0, fill = "#F4A582",
colour = NA, alpha = 0.5) +
  geom_line(data = dd4) + geom_ribbon(data = filter(dd4, x > qs4[[1]] & x < qs4[[3]]),
                                      aes(ymax = y), ymin = 0, fill = "#FDDBC7",
colour = NA, alpha = 0.5) +
  geom_line(data = dd5) + geom_ribbon(data = filter(dd5, x > qs5[[1]] & x < qs5[[3]]),
                                      aes(ymax = y), ymin = 0, fill = "#D1E5F0",
colour = NA, alpha = 0.5) +
    scale_y_continuous(limits = c(0,12), name = "Posterior Density") +
  scale_x_continuous(name ="Treatment vs Control",limits = c(-6,6) ) +
  annotate("text", x = c(qs1[[2]], qs2[[2]]-.1, qs3[[2]], qs4[[2]], qs5[[2]]-.2),
           y = c(6,12,11,12,8), label = c("AM","MY","WA","JS","WE"),
           parse = TRUE, size = 3) + theme_bw() + theme(axis.text = element_text(family = "Helvetica",
size = 18), text = element_text(family = "Helvetica", size = 12))
```

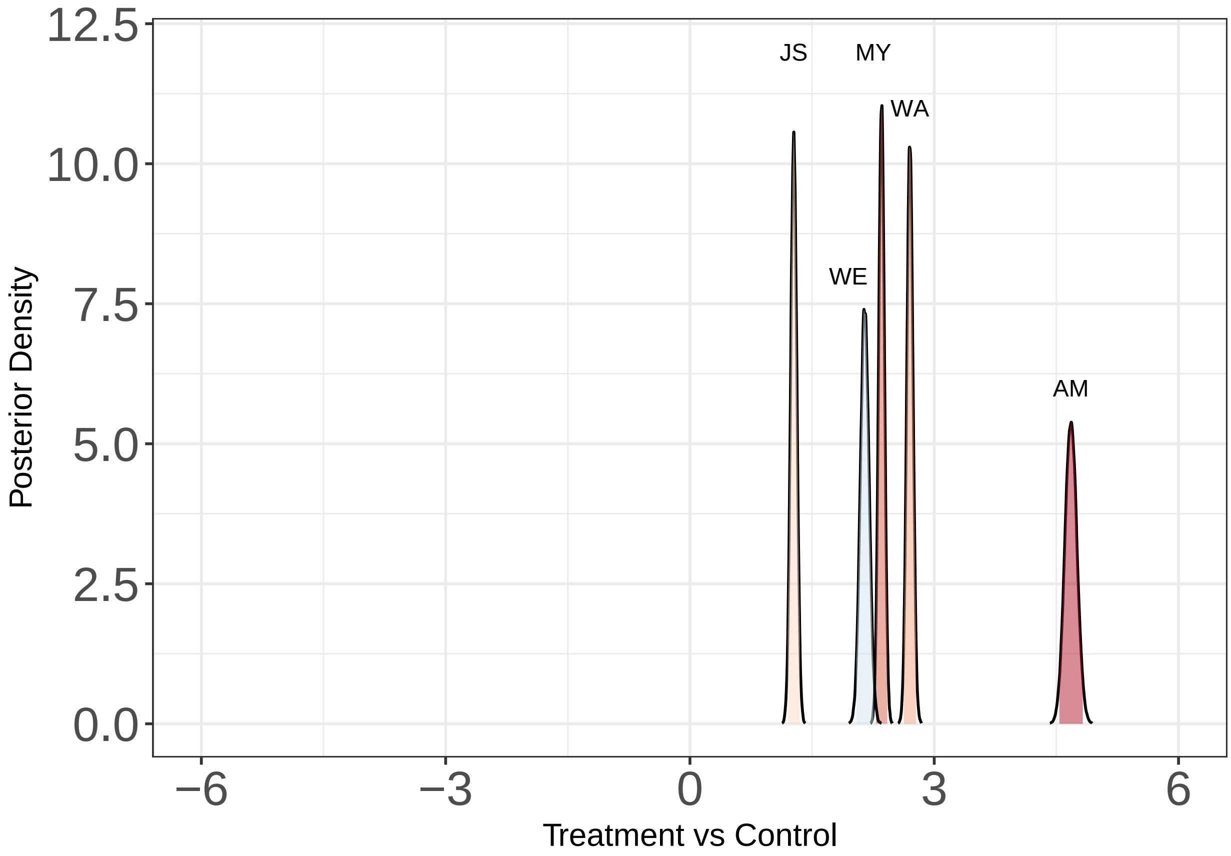


```r
overlap <- tibble(strain = c("AM","MY","WA","JS","WE"),
                  prob = c( ecdf( del1)(0),ecdf( del2)(0),ecdf( del3)(0),
                            ecdf( del4)(0),ecdf( del5)(0) ))

print(overlap)
```

```
## # A tibble: 5 x 2
##   strain  prob
##   <chr>  <dbl>
## 1 AM         0
## 2 MY         0
## 3 WA         0
## 4 JS         0
## 5 WE         0
```

Plots and analysis of the sporulation timing.

```r
#use the mean of the ancestral and the full posterior of the experimental
lq = 0.025
uq = 0.975
del1 <- (mean(samplesAncestral$AMAncTime) - samplesAM$AMCTiming)*(-1)
d1 <- density(del1)
dd1 <- with(d1, data.frame(x, y)) %>% filter(y > 0.01)
qs1 = quantile(ecdf( del1), prob = c(lq, 0.5, uq))
del2 <- (mean(samplesAncestral$MYAncTime) - samplesMY$MYCTiming)*(-1)
d2 <- density( del2)
dd2 <- with(d2, data.frame(x, y)) %>% filter(y > 0.01)
```

```
qs2 = quantile(ecdf(del2), prob = c(lq, 0.5, uq))
del3 <- (mean(samplesAncestral$WAAncTime) - samplesWA$WACTiming)*(-1)
d3 <- density( del3)
dd3 <- with(d3, data.frame(x, y)) %>% filter(y > 0.01)
qs3 = quantile(ecdf(del3), prob = c(lq, 0.5, uq))
del4 <- (mean(samplesAncestral$JSAncTime) - samplesJS$JSCTiming)*(-1)
d4 <- density( del4)
dd4 <- with(d4, data.frame(x, y)) %>% filter(y > 0.01)
qs4 = quantile(ecdf(del4), prob = c(lq, 0.5, uq))
del5 <- (mean(samplesAncestral$WEAncTime) - samplesWE$WECTiming)*(-1)
d5 <- density( del5)
dd5 <- with(d5, data.frame(x, y)) %>% filter(y > 0.01)
qs5 = quantile(ecdf(del5), prob = c(lq, 0.5, uq))



ggplot(data = dd1, aes(x, y)) +
  geom_line(data = dd1) + geom_ribbon(data = filter(dd1, x > qs1[[1]] & x < qs1[[3]]),
                                    aes(ymax = y), ymin = 0, fill = "#B2182B",
colour = NA, alpha = 0.5) +
  geom_line(data = dd2) + geom_ribbon(data = filter(dd2, x > qs2[[1]] & x < qs2[[3]]),
                                    aes(ymax = y), ymin = 0, fill = "#D6604D",
colour = NA, alpha = 0.5) +
    geom_line(data = dd3) + geom_ribbon(data = filter(dd3, x > qs3[[1]] & x < qs3[[3]]),
                                      aes(ymax = y), ymin = 0, fill = "#F4A582",
colour = NA, alpha = 0.5) +
  geom_line(data = dd4) + geom_ribbon(data = filter(dd4, x > qs4[[1]] & x < qs4[[3]]),
                                    aes(ymax = y), ymin = 0, fill = "#FDDBC7",
colour = NA, alpha = 0.5) +
  geom_line(data = dd5) + geom_ribbon(data = filter(dd5, x > qs5[[1]] & x < qs5[[3]]),
                                    aes(ymax = y), ymin = 0, fill = "#D1E5F0",
colour = NA, alpha = 0.5) +
    scale_y_continuous(limits = c(0,11), name = "Posterior Density") +
  scale_x_continuous(name ="Timing: Control vs Ancestral",limits = c(-2,2) ) +
  annotate("text", x = c(qs1[[2]], qs2[[2]], qs3[[2]], qs4[[2]], qs5[[2]]),
           y = c(10,10.3,10.6,10.9,6), label = c("AM","MY","WA","JS","WE"),
           parse = TRUE, size = 3) + theme_bw() + theme(axis.text = element_text(family = "Helvetica",
size = 18), text = element_text(family = "Helvetica", size = 12))
```

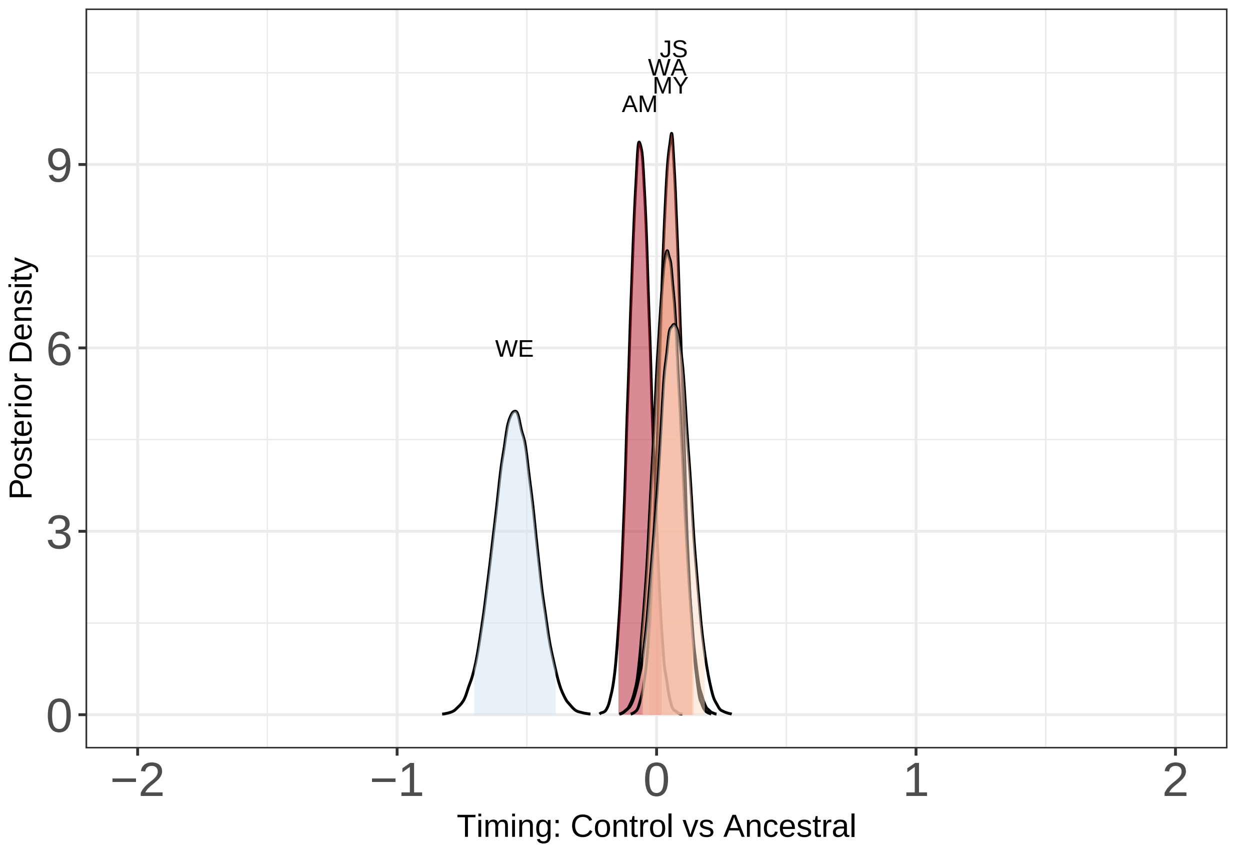


```
overlap <- tibble(strain = c("AM","MY","WA","JS","WE"),
                  prob = c( 1-ecdf( del1)(0),1-ecdf( del2)(0),
                        1-ecdf( del3)(0),1-ecdf( del4)(0),1-ecdf( del5)(0) ))

print(overlap)
```

```
## # A tibble: 5 x 2
##   strain   prob
##   <chr>   <dbl>
## 1 AM     0.0675
## 2 MY     0.900
## 3 WA     0.788
## 4 JS     0.862
## 5 WE     0
```

```
#use the mean of the ancestral and the full posterior of the experimental
lq = 0.025
uq = 0.975
del1 <- (mean(samplesAncestral$AMAncTime) - samplesAM$AMTTiming)*(-1)
d1 <- density(del1)
dd1 <- with(d1, data.frame(x, y)) %>% filter(y > 0.01)
qs1 = quantile(ecdf( del1), prob = c(lq, 0.5, uq))
del2 <- (mean(samplesAncestral$MYAncTime) - samplesMY$MYTTiming)*(-1)
d2 <- density( del2)
dd2 <- with(d2, data.frame(x, y)) %>% filter(y > 0.01)
qs2 = quantile(ecdf(del2), prob = c(lq, 0.5, uq))
del3 <- (mean(samplesAncestral$WAAncTime) - samplesWA$WATTiming)*(-1)
```

```
d3 <- density( del3)
dd3 <- with(d3, data.frame(x, y)) %>% filter(y > 0.01)
qs3 = quantile(ecdf(del3), prob = c(lq, 0.5, uq))
del4 <- (mean(samplesAncestral$JSAncTime) - samplesJS$JSTTiming)*(-1)
d4 <- density( del4)
dd4 <- with(d4, data.frame(x, y)) %>% filter(y > 0.01)
qs4 = quantile(ecdf(del4), prob = c(lq, 0.5, uq))
del5 <- (mean(samplesAncestral$WEAncTime) - samplesWE$WETTiming)*(-1)
d5 <- density( del5)
dd5 <- with(d5, data.frame(x, y)) %>% filter(y > 0.01)
qs5 = quantile(ecdf(del5), prob = c(lq, 0.5, uq))



ggplot(data = dd1, aes(x, y)) +
  geom_line(data = dd1) + geom_ribbon(data = filter(dd1, x > qs1[[1]] & x < qs1[[3]]),
                                      aes(ymax = y), ymin = 0, fill = "#B2182B",
colour = NA, alpha = 0.5) +
  geom_line(data = dd2) + geom_ribbon(data = filter(dd2, x > qs2[[1]] & x < qs2[[3]]),
                                      aes(ymax = y), ymin = 0, fill = "#D6604D",
colour = NA, alpha = 0.5) +
    geom_line(data = dd3) + geom_ribbon(data = filter(dd3, x > qs3[[1]] & x < qs3[[3]]),
                                        aes(ymax = y), ymin = 0, fill = "#F4A582",
colour = NA, alpha = 0.5) +
  geom_line(data = dd4) + geom_ribbon(data = filter(dd4, x > qs4[[1]] & x < qs4[[3]]),
                                      aes(ymax = y), ymin = 0, fill = "#FDDBC7",
colour = NA, alpha = 0.5) +
  geom_line(data = dd5) + geom_ribbon(data = filter(dd5, x > qs5[[1]] & x < qs5[[3]]),
                                      aes(ymax = y), ymin = 0, fill = "#D1E5F0",
colour = NA, alpha = 0.5) +
    scale_y_continuous(limits = c(0,11), name = "Posterior Density") +
  scale_x_continuous(name ="Timing: Treatment vs Ancestral",limits = c(-2,2) ) +
  annotate("text", x = c(qs1[[2]], qs2[[2]], qs3[[2]], qs4[[2]], qs5[[2]]),
           y = c(8,8.5,8.5,8,7.5), label = c("AM","MY","WA","JS","WE"),
           parse = TRUE, size = 3) + theme_bw() + theme(axis.text = element_text(family = "Helvetica",
size = 18), text = element_text(family = "Helvetica", size = 12))
```

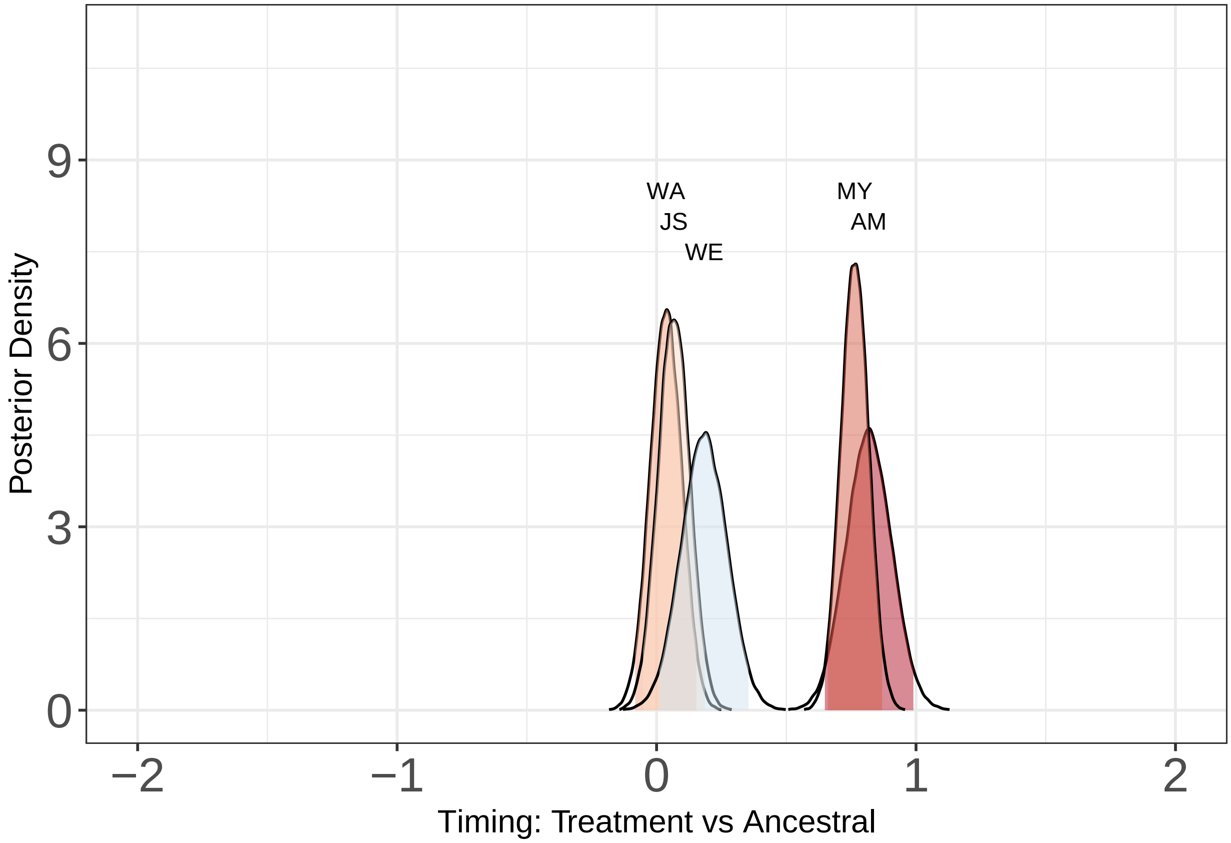


```
overlap <- tibble(strain = c("AM","MY","WA","JS","WE"),
                  prob = c( ecdf( del1)(0),ecdf( del2)(0),
                        ecdf( del3)(0),ecdf( del4)(0),ecdf( del5)(0) ))

print(overlap)
```

```
## # A tibble: 5 x 2
##   strain   prob
##   <chr>   <dbl>
## 1 AM     0
## 2 MY     0
## 3 WA     0.276
## 4 JS     0.138
## 5 WE     0.0197
```

```
lq = 0.025
uq = 0.975
del1 <- samplesAM$AMTTiming
d1 <- density(del1)
dd1 <- with(d1, data.frame(x, y)) %>% filter(y > 0.01)
qs1 = quantile(ecdf( del1), prob = c(lq, 0.5, uq))
del2 <- samplesMY$MYTTiming
d2 <- density( del2)
dd2 <- with(d2, data.frame(x, y)) %>% filter(y > 0.01)
qs2 = quantile(ecdf(del2), prob = c(lq, 0.5, uq))
del3 <- samplesWA$WATTiming
d3 <- density( del3)
```

```
dd3 <- with(d3, data.frame(x, y)) %>% filter(y > 0.01)
qs3 = quantile(ecdf(del3), prob = c(lq, 0.5, uq))
del4 <- samplesJS$JSTTiming
d4 <- density( del4)
dd4 <- with(d4, data.frame(x, y)) %>% filter(y > 0.01)
qs4 = quantile(ecdf(del4), prob = c(lq, 0.5, uq))
del5 <-samplesWE$WETTiming
d5 <- density( del5)
dd5 <- with(d5, data.frame(x, y)) %>% filter(y > 0.01)
qs5 = quantile(ecdf(del5), prob = c(lq, 0.5, uq))
```

```
ggplot(data = dd1, aes(x, y)) +
  geom_line(data = dd1) + geom_ribbon(data = filter(dd1, x > qs1[[1]] & x < qs1[[3]]),
                                      aes(ymax = y), ymin = 0, fill = "#B2182B",
colour = NA, alpha = 0.5) +
  geom_line(data = dd2) + geom_ribbon(data = filter(dd2, x > qs2[[1]] & x < qs2[[3]]),
                                      aes(ymax = y), ymin = 0, fill = "#D6604D",
colour = NA, alpha = 0.5) +
    geom_line(data = dd3) + geom_ribbon(data = filter(dd3, x > qs3[[1]] & x < qs3[[3]]),
                                        aes(ymax = y), ymin = 0, fill = "#F4A582",
colour = NA, alpha = 0.5) +
  geom_line(data = dd4) + geom_ribbon(data = filter(dd4, x > qs4[[1]] & x < qs4[[3]]),
                                      aes(ymax = y), ymin = 0, fill = "#FDDBC7",
colour = NA, alpha = 0.5) +
  geom_line(data = dd5) + geom_ribbon(data = filter(dd5, x > qs5[[1]] & x < qs5[[3]]),
                                      aes(ymax = y), ymin = 0, fill = "#D1E5F0",
colour = NA, alpha = 0.5) +
    scale_y_continuous(limits = c(0,11), name = "Posterior Density") +
  scale_x_continuous(name ="Timing: Treatment vs Control",limits = c(-2,2) , breaks = c(-2,-1,0,1,2)) +
  annotate("text", x = c(qs1[[2]], qs2[[2]], qs3[[2]], qs4[[2]], qs5[[2]]+.1),
           y = c(5.5,8,7.3,7.3,5), label = c("AM","MY","WA","JS","WE"),
           parse = TRUE, size = 3) + theme_bw() + theme(axis.text = element_text(family = "Helvetica",
size = 18), text = element_text(family = "Helvetica", size = 12))
```

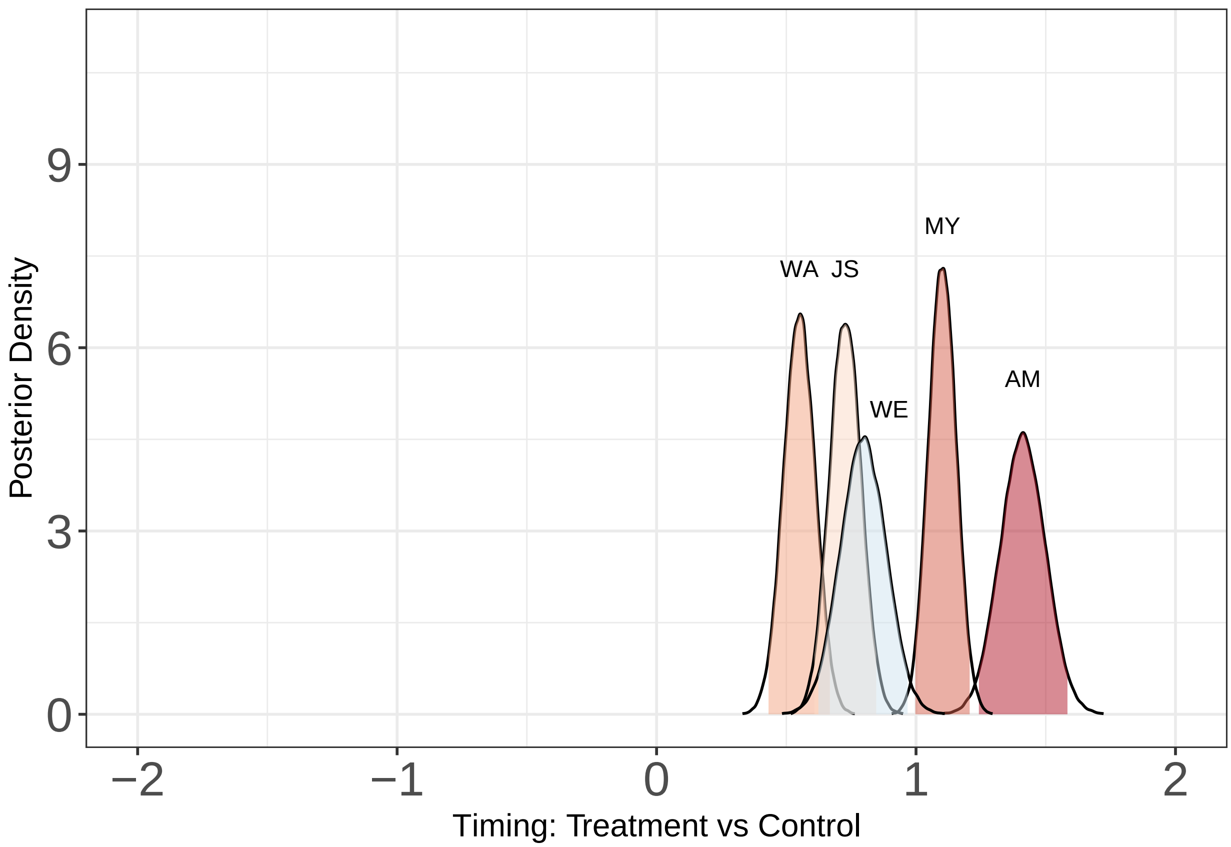


```
overlap <- tibble(strain = c("AM","MY","WA","JS","WE"),
                  prob = c( ecdf( del1)(0),ecdf( del2)(0),ecdf( del3)(0),
                      ecdf( del4)(0),ecdf( del5)(0) ))

print(overlap)
```

```
## # A tibble: 5 x 2
##   strain  prob
##   <chr>  <dbl>
## 1 AM         0
## 2 MY         0
## 3 WA         0
## 4 JS         0
## 5 WE         0
```
